# The balance of interaction types determines the assembly and stability of ecological communities

**DOI:** 10.1101/643478

**Authors:** Jimmy J. Qian, Erol Akçay

**Affiliations:** Department of Biology, University of Pennsylvania 433 S University Ave, Philadelphia, PA 19104, USA

## Abstract

What determines the assembly and stability of complex communities is a central question in ecology. Past work has suggested that mutualistic interactions are inherently destabilizing. However, this conclusion relies on assuming that benefits from mutualisms never stop increasing. Furthermore, almost all theoretical work focuses on the internal (asymptotic) stability of communities assembled all-at-once. Here, we present a model with saturating benefits from mutualisms and sequentially assembled communities. We show that such communities are internally stable for any level of diversity and any combination of species interaction types. External stability, or resistance to invasion, is thus an important but overlooked measure of stability. We demonstrate that the balance of different interaction types governs community dynamics. Mutualisms may increase external stability and diversity of communities as well as species persistence, depending on how benefits saturate. Ecological selection increases the prevalence of mutualisms, and limits on biodiversity emerge from species interactions. Our results help resolve longstanding debates on the stability, saturation, and diversity of communities.

## Introduction

An enduring central question of ecology is what makes ecological communities stable. Elton [1] famously argued that complex communities should be more resistant to invasion by new species. By the 1970s, this idea was established as a cornerstone of ecology [2]. However, the field was given a jolt by the seminal theoretical work of May who showed that complex ecological communities in fact tend to be less stable, in the sense of returning to equilibria from small perturbations to existing species [3, 4]. This set off the great complexity-stability debate, as nature abounds with complex communities containing large numbers of species. A large theoretical and empirical literature developed studying different notions of complexity, diversity, and stability [2, 5–9]. Yet despite this enormous attention, theoretical work has left important aspects of the question understudied.

The first understudied aspect is accounting for diversity of interaction types between species (e.g., mutualism, competition, and exploitation), which is expected to affect stability and assembly of ecological communities. May’s original analyses considered random interactions among species, while subsequent studies initially considered species networks connected only by a single interaction type, focusing on purely competitive, exploitative (predator-prey), or mutualistic communities. Only more recently has theoretical work started to incorporate multiple interaction types and asked how the balance of interaction types within a community affects its stability [10–18]. Most of these studies find that mutualistic interactions have destabilizing effects on communities, which are only stable when mutualistic interaction strengths are weak or asymmetric [13–16]. These results would appear to lead to the conclusion that mutualisms must play relatively small roles in complex communities.

However, this conclusion rests largely on neglect of a second important aspect of the complexity-stability question: the role of nonlinear functional responses. It is easy to see that the destabilizing effect of mutualisms stems from the fact that unbounded positive feedbacks can arise from Lotka-Volterra models with a linear functional response, where the per capita effect of one species on another is independent of their population sizes. But linear functional responses are biologically unrealistic when studying mutualisms [19–24]. The benefits a focal species receives per capita from a mutualist partner cannot increase without limit as the partner species’ population size increases; they must saturate eventually [20], as the focal species become limited by resources or services other than the one the mutualistic partner is providing. Therefore, nonlinear functional responses with saturating effects for mutualistic interactions are more realistic models to consider the effects of mutualisms on community stability [20–24]. Yet they remain critically underused, having been used to model mutualistic communities only in a handful of studies [10, 11, 22, 25–28].

Furthermore, exploitative interactions are also expected to exhibit nonlinear saturating functional responses [24, 29–31], since predators can become satiated with high levels of prey (either literally, i.e., eating to their physiological limit, or time budget-wise, due to the handling time for each prey). Basic theory shows that the effects of saturating functional responses in pairwise mutualistic and exploitative interactions should be opposite, stabilizing in the former case, while destabilizing in the latter. Thus, the overall effect of incorporating saturating responses on community stability likely depends on the balance of different kinds of interactions.

Finally, most of the theory on the relationship between diversity and stability is concerned with the internal stability of communities, i.e., whether a collection of species that coexist at an equilibrium will return to this equilibrium if perturbed (also called asymptotic stability). Theoretical models typically approach this question with random communities assembled all at once, where an interaction matrix of a certain size is populated from some distribution, and its stability determined. However, communities in nature assemble sequentially, as new species enter and existing ones go extinct. This means that community matrices drawn all at once may not be representative of natural communities, and that external stability of communities, i.e., whether new species can invade (and conversely, existing species go extinct), will play an important role in determining the structure of communities. With few exceptions [32, 33], the relationship between external stability and diversity has been severely understudied relative to internal stability.

Here, we present an approach that jointly addresses these three neglected aspects of the diversity-stability debate. We construct a model of community assembly where communities emerge from sequential invasion of randomly selected species, rather than studying community matrices assembled all at once. We vary the proportions of competitive, exploitative, and mutualistic interactions in the invader pool to ask how the balance of interaction types affects the structure and stability of assembled communities. We use saturating functional responses for mutualistic and exploitative interactions. We show that in this setting, the problem of internal stability effectively vanishes, while the problem of external stability – Elton’s original question – comes into focus.

## Results

We first give an overview of our model (see Methods for detailed model description). We model sequentially assembled communities of species that can have competitive, mutualistic, or exploitative interactions with each other. We use saturating (Holling Type II) functional responses [20, 29] with half-saturation constant *h* for mutualistic and exploitative interactions. This setting raises a question that does not come up with linear functional responses: whether each mutualistic or exploitative interaction is unique, or whether they are interchangeable with each other. For mutualisms, if each partner provides a unique type of benefit (different nutrients or services) to a focal species, the interaction strength should saturate in that partner’s density only. We call this scenario the Unique Interactions Model (UIM). At the other extreme, if all mutualisms provide the same type of benefit, then the functional responses should saturate in the total density of all mutualist partners. We call this scenario the Interchangeable Interactions Model (IIM). Similar considerations apply to exploitative interactions, where prey and predators may be unique or interchangeable with each other. Both types of models have been considered in previous studies [10, 28], though their results have not been directly compared to one another. We would expect interactions in nature to be at some intermediate between the completely unique and completely interchangeable models. Thus, we provide results for both the UIM and IIM, focusing on UIM to introduce our main conclusions, and later comparing the results with the IIM. We assume that competitive interactions retain the oft-assumed linear (Type I) functional response, and hence that the issue of interchangeability does not arise for competitive interactions.

Each community starts with a small number of species and grows by the repeated invasion of new species. Each time a new species arrives in the community, its interactions with existing species are generated randomly, with average connectivity (i.e., probability of any interaction) *C*, and frequencies *P_m_*, *P_c_*, *P_e_* for mutualistic, competitive, and exploitative interactions, respectively. When we introduce a new species, it can either have a positive growth rate when rare and is able to invade, in which case it is added into the community, or it will have a negative growth rate when rare, and thus will not be added to the community. In the former case, we simulate the dynamics of the community until it settles to an ecological steady state, which might be an equilibrium or a limit cycle. We introduce a new species to this steady state, reflecting a separation of timescales assumption [34–36]. At each equilibrium, we generate new species until an invasion occurs and track the probability of invasion. Species can also go extinct. We repeat this process until the community assembly process reaches a steady state in species richness, quantified using a unit root test. In the main text, we present results setting connectivity *C* = 0.5, but all our findings are robust across different levels of connectivity and also different values of *h* (Supplementary Information).

In our model, new species are introduced into communities without directly controlling the species richness. In other words, the biodiversity of a community is constrained by emergent limits on community complexity. We find that the relative proportions of interaction types govern the emergent species diversity in assembled communities (Fig. 1). Notably, as long as communities have nonzero competition, they saturate as large numbers of invasions occur and the species richness converges to some steady state (Fig. 1a). The steady state is characterized by small stochastic oscillations around some baseline value, so that a sequence of consecutive invasions is offset by a subsequent cascade of extinctions, even when thousands of invasions occur. This holds for both the unique and interchangeable interactions models. In both models, communities with more mutualism and less competition show higher diversity at this steady state (Fig. 1a,c, Fig. 6a), with the mutualism effect more pronounced in the UIM, and competition effect in the IIM. Regardless of *C* or *h*, community diversity saturates (Supplementary Figs. 2,3 for UIM; 19, 20 for IIM). The lower the connectivity, the higher the steady state species richness, but the balance of interaction types yields the similar qualitative patterns patterns regardless of connectivity and saturation constant.

**Figure 1:**
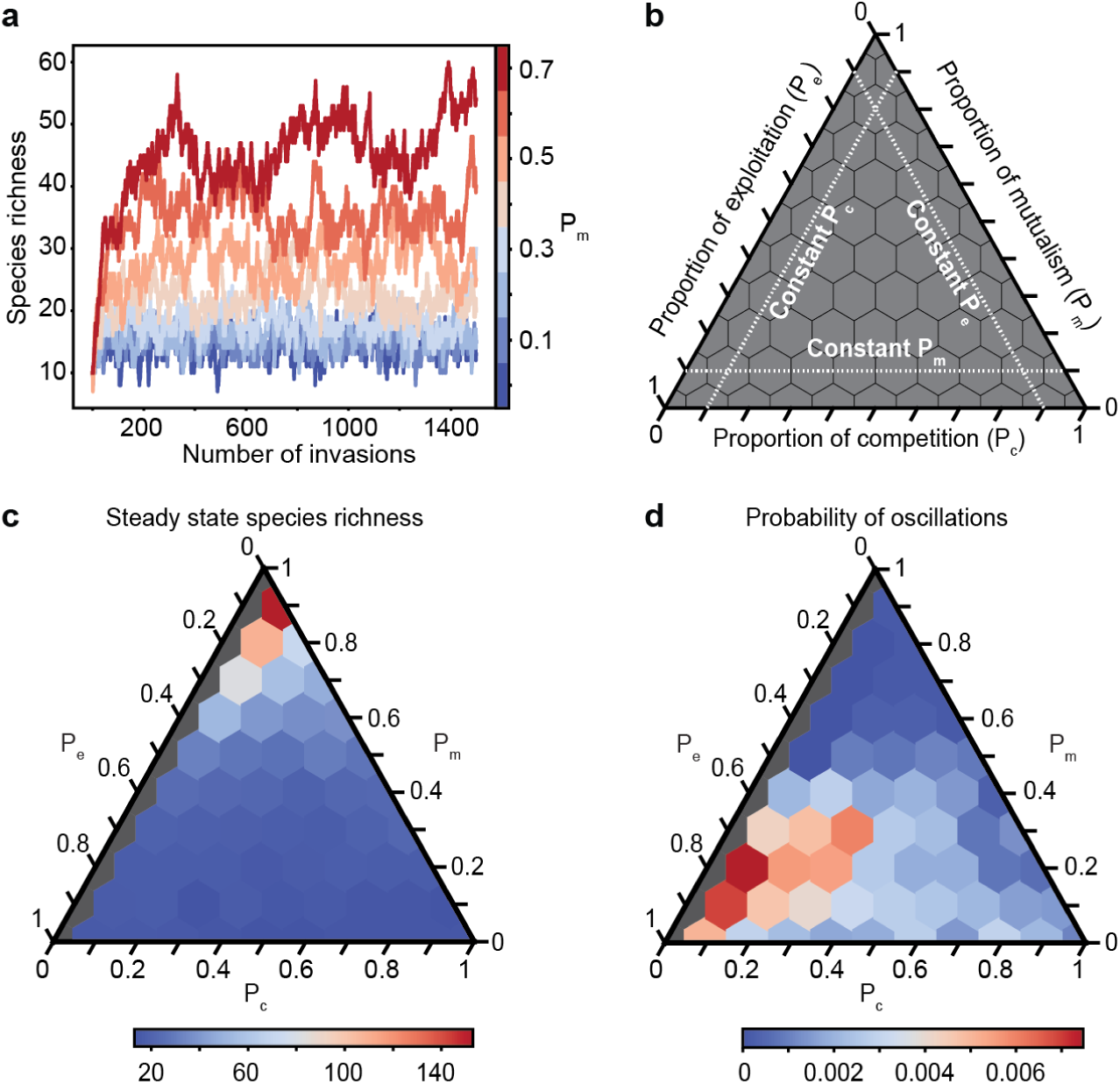
The balance of interaction types determines the species richness at which communities saturate under the unique interactions model (UIM). (a) Species richness in representative communities with *P_c_* = 0.3 and varying levels of *P_m_*, represented by the color. Each line corresponds to a community over 1500 species invasions that occur at equilibria. The same trend is observed for all *P_c_ >* 0. (b) A schematic on how to read ternary plots [37]. The three lines show directions in which each parameter is constant. These are in the same directions as the corresponding axes tick marks. Each hexagon represents a single combination of the three interaction types, given by the three axes. (c) A ternary plot showing the steady state species richness of different communities with varying proportions of competition, mutualism, and exploitation. The color represents the species richness. Gray communities have no competition and are not considered since they are biologically unrealistic. (d) Probability that an equilibrium in the community history shows oscillations in population dynamics. This is defined as the number of times oscillations occur divided by the total number of equilibria. The color represents the probability.

Asymptotic stability provides information on whether communities return to an equilibrium state after small perturbations [3]. In our simulations, the population dynamics almost always converge to an equilibrium, in rare cases settling to stable oscillations (Supplementary Fig. 4). We never observe populations grow boundless, due to the saturating functional responses for mutualistic and exploitative interactions as well as intraspecific density dependence. The probability of oscillations is very low for all proportions of interaction types for both IIM and UIM (Fig. 1d, Supplementary Fig. 21), and this holds for almost all combinations of *C* and *h* (Supplementary Figs. 5, 21). In particular, we find that mutualistic interactions do not result in asymptotically unstable communities even when they are strong and abundant, consistent with past results in purely mutualistic communities with saturating functional responses [22]. Instead, internal stability is a ubiquitous property of sequentially assembled communities with saturating functional responses in mutualistic and exploitative interactions.

On the other hand, we find that the external stability, or the resistance of communities to invasion, is a critical determinant of community structure and function. Even though this was the original question motivating Elton [1] and many authors noted invasibility can be a measure of community instability [2, 8, 9, 38, 39], resistance to invasion from the outside has largely been overlooked in the theoretical debate on the complexity-stability relationship.

The species richness of communities is mediated through balance of invasion and extinction rates, which in turn depend on the balance of interaction types. Fig. 2a,b shows that in the UIM, the probability a new species can invade decreases with high proportion of competitive interactions, as one might expect. However, it also shows that, contrary to what one might expect, higher proportion of mutualisms also leads to lower probability of invasion. This is due to the fact that highly mutualistic communities allow resident species to achieve high population sizes, such that an invader with a competitive interaction with a resident will face strong competition. Because the benefits from mutualistic interactions saturate but the cost of competitive interactions does not, even a few competitive interactions can be enough to keep an invader out. Thus, mutualisms with saturating functional responses increase the competitive burden to invaders and render communities externally stable.

**Figure 2:**
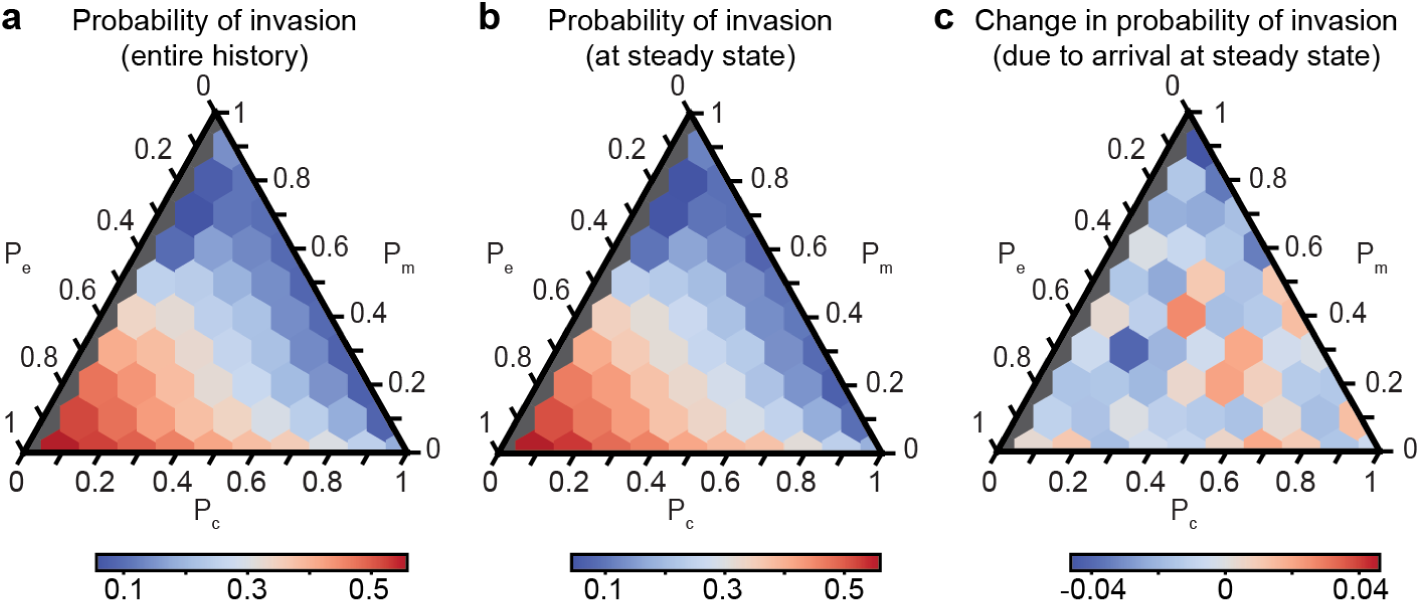
External stability is determined by the proportion of interaction type in the unique interactions model (UIM). (a) External stability throughout the entire community history. Colors indicate the probability of invasion. Gray communities have no competition and are not considered. (b) External stability after species richness has converged to the steady state. Colors indicate the probability of invasion. (c) The difference in probability of invasion between the transient state and the steady state. The colors represent the change in probability.

More specifically, for communities with moderate proportion of exploitative interactions, increasing mutualism while decreasing competition monotonically enhances external stability. At low proportions of exploitative interactions, external stability is maximized at intermediate levels of mutualism. Conversely, holding competition constant, external stability increases with increasing mutualism (except when *P_c_* is very low). For low and moderate proportions of mutualistic interactions, increasing competition (and thereby decreasing exploitation) augments stability, whereas when *P_m_* is high, decreasing competition increases stability. These trends are observed both throughout the entire community history (Fig. 2a) and during the steady state after diversity has converged (Fig. 2b). A community’s stability does not change as it leaves its transient phase (before species diversity converges) for the steady state (Fig. 2c). Any changes in probability of invasion are small, and there is no clear pattern throughout the different communities. These trends are qualitatively robust for all *C* and *h* (Supplementary Figs. 6, 7, 8).

In contrast to the UIM, mutualisms do not markedly increase the external stability of communities in the IIM (Fig. 6a, Supplementary Fig. 22). Under the IIM, invasion probabilities are generally low and show no strong pattern with the proportion of mutualistic interactions and weakly decrease with exploitative interactions. Looking at the mean population sizes in the IIM confirms that this contrasting result is due to the fact that varying the proportion of mutualisms and exploitative interactions in the IIM does not cause population sizes to change significantly (Fig. 6d). Thus, the mechanism that stabilizes communities against invasion in the UIM is unavailable in the IIM. This shows that mutualisms can be externally stabilizing as long as they allow resident species to achieve high densities.

Our model also illustrates how species interactions can modulate the relationship between species richness and resistance to invasion in different ways. Under the UIM, we can observe both a positive and negative relationship, depending on which interaction proportions are varied: holding the proportion of mutualistic interactions constant and trading off exploitative against competitive interactions, species richness and invasion probability are positively correlated, producing what has been termed the “invasion paradox” [40] (Fig. 1c; Fig. 2a,b; Supplementary Figs. 3, 6, 7). However, keeping the proportion of competitive interactions constant and trading off mutualisms against exploitative interactions, we get a negative relationship between species richness and invasion probability. Here, more mutualistic communities have more species but are less likely to be invaded. In contrast, under the IIM, the relation between invasion probability and species richness is largely negative, and modulated by the proportion of competitive interactions (Fig. 6b,c).

The other determinant of community richness is the probability an existing species goes extinct. Our model shows that this probability also depends on the balance of interaction types. We define the probability of extinction simply as the number of species that went extinct divided by the total of number of species that existed during the relevant part of the community history. In the UIM, for any constant level of competition, decreasing exploitation and increasing mutualism decreases the probability of extinction (Fig. 3a,b; Supplementary Figs. 9, 10). For constant moderate or high exploitation, extinction is minimized for low competition and high mutualism (for high *h*, this is true for all levels of exploitation). With constant mutualism, increasing exploitation or decreasing competition decreases extinction. As expected, these trends are the opposite of those seen in Fig. 1c for steady state species richness. In the IIM, extinction probability again is mostly a function of the fraction of competitive interactions (Fig. 6c, Supplementary Fig. 25), without very strong dependence on the fraction of exploitative or mutualistic interactions.

**Figure 3:**
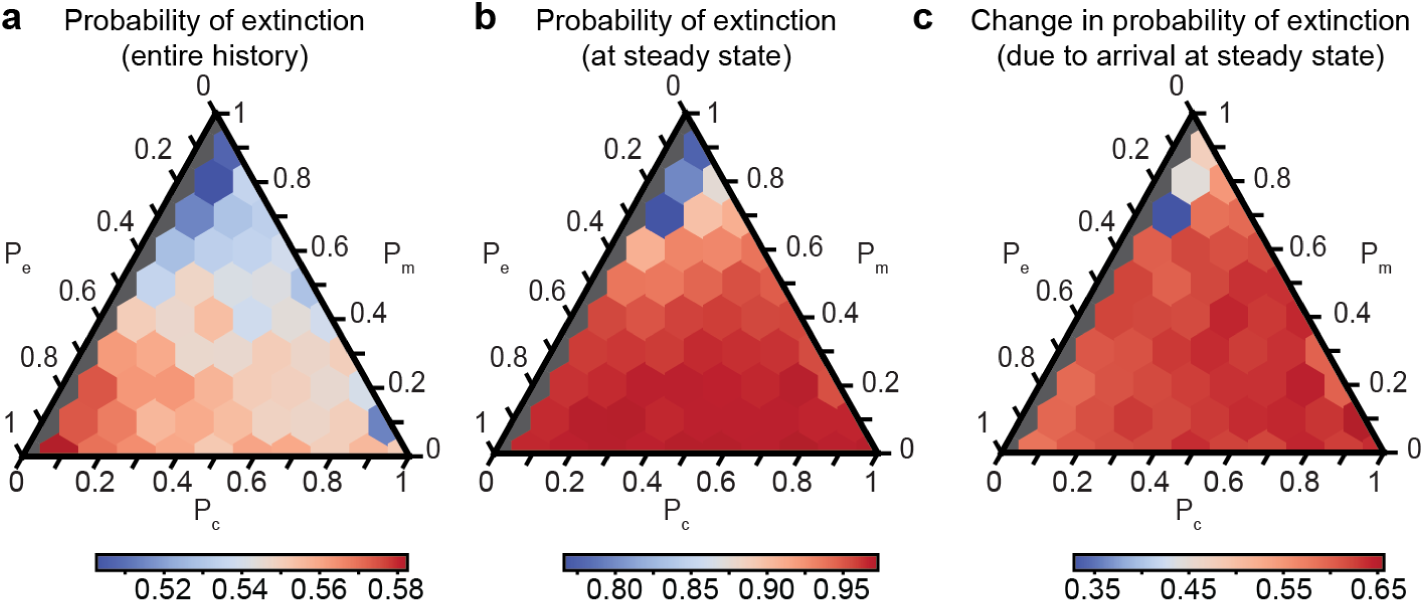
The probability of extinction is determined by the proportion of interaction types in the unique interactions model. (a) Extinction throughout the entire community history. Colors indicate the probability of extinction; gray communities have no competition and are not considered. (b) Probability of extinction after species richness has converged to the steady state. Colors indicate the probability of extinction. (c) The difference in probability of extinction between the transient state and the steady state. The colors represent the change in probability.

In both the UIM and IIM, community saturation and emergence of a limit on biodiversity occur due to changes in extinction probability as opposed to invasion, which stays constant before and after convergence of species diversity to the steady state (Fig. 2c; Supplementary Figs. 8, 24), whereas the probability of extinction increases after the steady state is reached (Fig. 3c; Supplementary Figs. 11, 27). The steady state species richness is maintained when invasion of new species precipitates the extinction of older species, rather than through a resistance to invasion by new species. As a result, the distribution of species persistence is skewed right for all communities, and most species do not survive very long (Fig. 4a; Supplementary Fig. 12). To quantify how persistence varies with species interactions, we define relative persistence as the number of equilibria that the species remains extant normalized to the number of equilibria in the respective simulation. In the UIM, high proportion of mutualisms and low proportions of exploitative interactions both favor slow species turnover and longer species persistence (Fig. 4; Supplementary Figs. 13,14) as they both reduce the invasion and extinction probabilities. In particular, keeping the level of competition constant and increasing more mutualistic interactions lengthens the tail of the persistence distribution so that some species survive much longer than others (Fig. 4a). In contrast, species persistence in the IIM again mostly depends on the proportion of competitive interactions (Supplementary Figs. 28, 29), which modulates both invasion and extinction probabilities.

**Figure 4:**
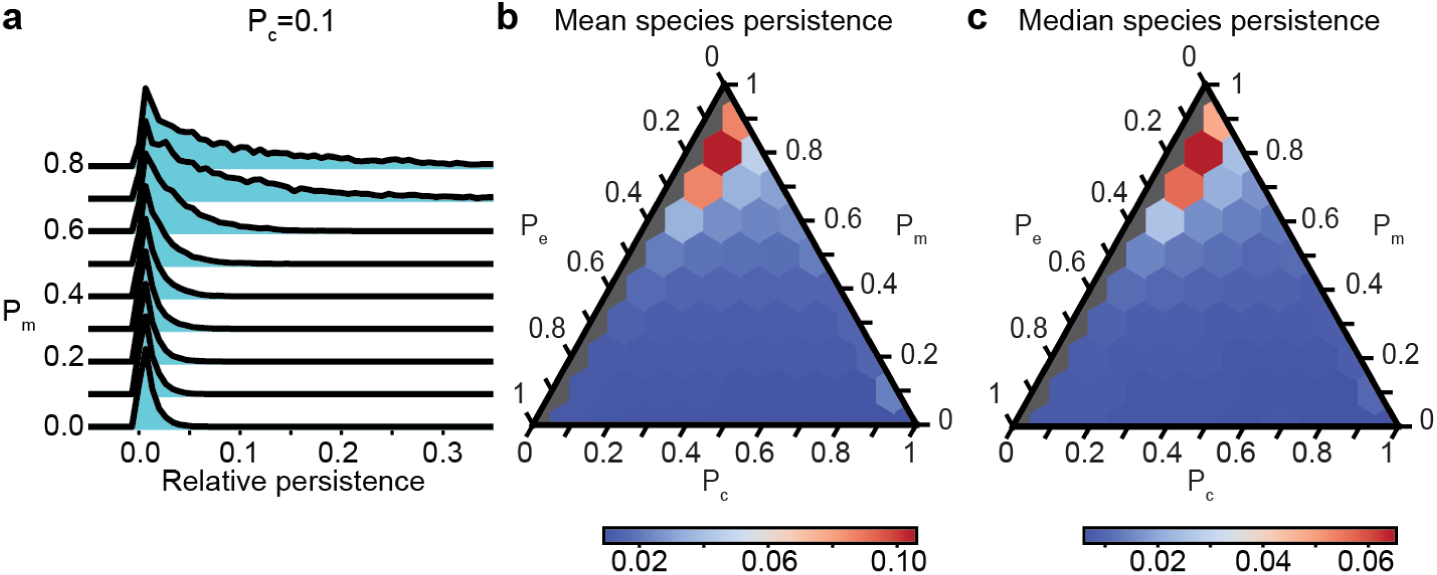
Species persistence in the unique interactions model is determined by the proportion of interaction types. (a) Normalized histograms of the relative persistence of species in communities with *P_c_* = 0.1 and varying *P_m_*. This same trend is observed for all *P_c_*. (b) The mean relative persistence of all species that existed within a community. The color represents the mean relative persistence. (c) The median relative persistence of all species that existed within a community. The color represents the median relative persistence.

Finally, we consider ecological selection (or filtering) of interaction types during community assembly. Our model throws at communities randomly generated species that have mutualistic, competitive, and exploitative interactions at proportions *P_m_*, *P_c_*, and *P_e_*, respectively (we call this the extrinsic distribution). Although candidate invader species are generated randomly (see Methods), the ones that are able to invade and persist in a community may be biased towards one or the other interaction type. Indeed, we observe the signature of such selection acting on the distribution of interaction types in a community in both the UIM and IIM. We find that selection acts to decrease competition, increase mutualism, and slightly increase exploitation, with a more dramatic increase in exploitation at lower levels of mutualism (Fig. 5; Supplementary Fig. 30). This selection is caused by a combination of biases in the invasion and extinction rates. Successful invaders generally show the same bias for more mutualism and less competition (Fig. 5; Supplementary Fig. 31). This is not surprising since an invader that benefits from existing species will have an easier time invading than one that suffers competition from them. One exception is in the UIM, when exploitation is high and competition is low, where invaders tend have more competition. Extinction generally is biased in the same way as the overall community composition (Fig. 5c; Supplementary Fig. 32), except again in the high-exploitation, low-competition cases in the UIM. The difference in interaction types between species that invade and those that go extinct is generally small (Fig. 5d; Supplementary Fig. 33), except again in the UIM for high-exploitation, low-competition communities, where extinction is biased towards lower competition. The finding that ecological selection increases the proportion of mutualisms is generally robust to variation in connectivity or the half-saturation constant (Supplementary Figs. 15, 16, 17, 18, 30, 31, 32, 33), though the latter can have slightly differing effects on other interaction types.

**Figure 5:**
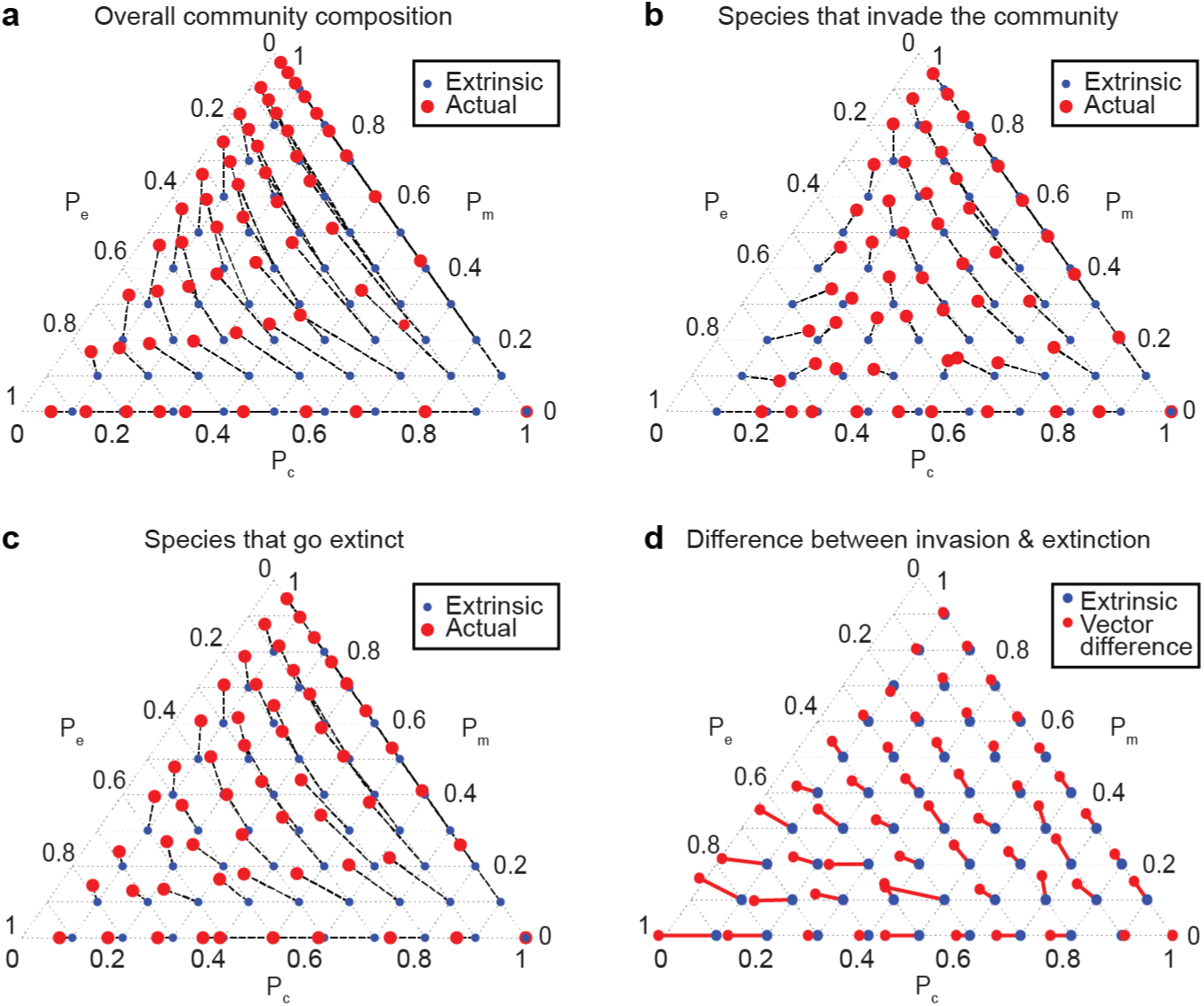
Selection acts to increase mutualism, and sometimes increase exploitation and decrease competition in the UIM. (a) The extrinsic (blue points) and the actual (red circles) interaction proportions of all species within each community. Dotted lines connect points from the same community, and can be interpreted as the direction of selection. See Methods for more details. (b) The extrinsic (blue points) and the actual (red circles) interaction proportions of species that successfully invade each community. Dotted lines connect points from the same community, and can be interpreted as the direction of selection. (c) The extrinsic (blue points) and the actual (red points) interaction proportions of species that go extinct. Dotted lines connect points from the same community. (d) The vector difference between species that go extinct and species that invade the community are plotted as red lines with red points at the end. The vectors start from the extrinsic proportions (blue points) and are defined as the interaction types of species that go extinct minus the interaction types of invaders.

## Discussion

Our model incorporates two key biologically realistic assumptions: saturating functional responses for mutualistic and exploitative interactions, and sequential selection of candidate invader species during community assembly. We show that under these assumptions, internal stability of communities is always guaranteed (in the form of either equilibria or stable limit cycles) with all combinations of interaction types, species diversity, and community connectivity. Part of this result can be explained by the fact that, by construction (and for biological realism), we restrict our attention to feasible communities of species that actually have positive abundance at equilibria. An early but relatively overlooked result by Roberts [41] shows that feasible equilibria tend to also be internally stable. Consistent with this, Stone [42, 43] showed that as variability of interaction strengths increases, communities become unfeasible (some species go to negative abundance at equilibrium) sooner than they become asymptotically unstable, and that in a broad range of cases, feasibility implies asymptotic stability. Thus, feasible communities tend to be stable. In addition, we find that mutualistic interactions with saturating functional responses, even if saturation only happens at high population densities, promote internal stability also consistent with previous results [28]. Overall, our results imply that internal stability is a ubiquitous property of communities regardless of their complexity and composition of species interactions, and therefore unlikely to be a differentiating feature of natural communities.

On the other hand, external stability, i.e., the ability of a community to resist invasion, emerges in our model as a crucial factor that varies with community structure and is pivotal in determining species richess. Yet, external stability has received much less attention than internal stability [1, 8, 9, 32, 33, 38, 39]. A notable exception is a model by Law and Morton [33] where they consider the sequential assembly of a purely exploitative community (i.e., a food web) from a finite species pool. By considering the endogenous assembly of communities from a large source pool with different types of interactions, we are able to answer a major outstanding question of ecology: how does the balance of species interactions drive community assembly and structure? Our model reveals how the external stability of communities, their steady state richness, and the extinction and persistence of organisms are determined by the balance of interaction types.

Our results show that mutualisms can play a key role in determining the assembly and stability of communities. In the unique interactions model, we find that when holding either competition or exploitation constant, mutualism augments internal stability, decreases extinction, increases species persistence, and thereby decreases species turnover. Interestingly, in the interchangeable interactions model, where mutualistic (and exploitative) interactions saturate in the total density of all interaction partners, these effects are absent or weaker. In particular, mutualisms no longer clearly increase external stability in the IIM. This is adding more mutualistic interactions in the IIM does not actually lead to generation of more gross mutualistic benefit in the community as most functional responses tend to be in the saturated range already, and adding new mutualistic partners do not noticeably increase the mutualistic benefit a focal species gets. Thus, more mutualistic interactions in the IIM do not lead to (much) higher population sizes for resident species, which takes away the mechanism by which mutualisms stabilize communities against external invaders in the UIM. In nature, interactions are likely to be of some intermediate type between completely unique and interchangeable, and thus might increase resident species densities to different degrees. Our model therefore predicts that to the extent that having more mutualistic interactions increase densities, they will also render communities more externally stable.

We find that mutualisms also raise the emergent limits on diversity, with a stronger effect in the UIM, allowing the community to saturate at a higher species richness. Accordingly, selection acts to increase mutualism during community assembly. Our demonstration that the ratio of mutualism to competition is an important determinant of community diversity, with a high ratio leading to high diversity, is consistent with a previous empirically derived model [17]. However, our findings contradict generalizations of May’s original stability-complexity results [3], which find not only that mutualism is destabilizing, but that it is even more destabilizing as the product of species richness and connectivity increases [13, 14, 44]. Such a destabilizing effect of mutualism contradicts empirical observations of frequent mutualistic interactions in communities with high diversity [44, 45]. In our model, the internal stability is not compromised with increasing species richness since sequentially assembled communities are selected for feasibility and the benefits from mutualisms saturate at high densities, both realistic features of communities in nature. The first feature means that communities can only get richer if they remain feasible, and therefore no species blows up in density, which would cause some other species to go extinct. The second means that the blowing up in density is unlikely in the first place, as the positive feedback between mutualist densities is cut short at high enough densities.

Previous work by Mougi and Kondoh [10–12] also showed that mutualisms may not be inherently destabilizing, but they focused exclusively on internal stability of fixed size communities. Furthermore, they assume that the total interaction strength of each species for each interaction type is constrained. This means that as mutualisms become common each mutualistic link gets weaker, in addition to interactions saturating in total species abundance (similar to our IIM) in their saturating functional response model. In contrast, Kawatsu and Kondoh [28] considered a case that is closer to our UIM, and showed that saturating functional responses can stabilize fixed size communities as in our model. We add to these results by showing that saturating responses with ecological selection is enough to almost guarantee internal stability in both the in the unique and interchangeable interaction models. Another recent line of work by Butler and O’Dwyer [46, 47] explicitly modeled species limited by resources that can be produced by other species. This model allows saturating functional responses in mutualisms to emerge endogenously from resource limitation. Consistent with our results, they found that feasible mutualistic communities are stable even if mutualistic links are strong, provided the exchange of resource is reciprocal between species. More generally, resource-mediated interactions tend to produce saturating functional responses as opposed to linear [48]. An important future direction is to model the external stability of communities with resource-mediated interactions.

One of our main findings is that all of the communities in our model show an emergent limit on species richness. This is true regardless of the connectivity or distribution of interaction types (as long as there is nonzero competition, which is certainly true in natural communities) or whether interactions are unique or interchangeable. In other words, all communities become saturated when mutualism and exploitation have saturating effects and communities are sequentially assembled. This result contributes to an ongoing debate that has centered around the relative importance of local biotic interactions and regional processes in determining community richness and saturation [49–54]. We show that the composition of local species interactions can generate an intrinsic limit on community diversity. The exact effect of the balance of interactions on species richness depends somewhat on whether interactions are unique or interchangeable but diversity generally is higher with lower competition and higher mutualisms. This saturation comes about even though we effectively model an infinite species pool with interactions randomly drawn according to the desired proportions, rather than a finite regional or metacommunity pool. Regardless of how many species attempt to invade the community, the species richness fluctuates around some steady state value that depends only on the distribution of interaction types. Our results thus suggest that species interactions are sufficient for community saturation on ecological timescales.

Our results on species richness and its equilibration add a new dimension to MacArthur and Wilson’s classical theory of island biogeography [55], which states that the species richness of an island is controlled by immigration and extinction patterns. We show that these patterns are in turn governed by the balance of species interactions. We therefore posit that the distribution of interaction types is a fundamental determinant of island biodiversity, and of community assembly in general. MacArthur and Wilson investigated how an island’s area and distance to the mainland affect the rates of immigration and extinction, and predicted an equilibrium of species diversity when these two rates are equal. We extend their analysis by considering how species interact with one another. We show that all communities can exhibit a dynamic equilibrium of species richness. Their equilibration model requires the extinction rate to increase with species diversity, and we provide insight into the underlying proportions of interactions that allow this to happen. In fact, we reveal that an increasing extinction rate (rather than a decrease in invasion rate) is indeed what drives the convergence of diversity. Our findings that species interactions can explain the distribution of island species is consistent with previous results [32].

The patterns we uncover also shed new light on the relationship between diversity and invasibility of a community, where evidence for both a negative and positive relationship exists [40]. Elton’s hypothesis that diversity decreases invasibility implicitly assumes that competition is the primary driver of community composition and invasibility. This reasoning was confirmed by Case [32], who modeled invasion of new species and extinction of resident ones in purely competitive communities. We show that accounting for different interaction types may help explain the invasion paradox. Rather than a direct correlation between resistance to invasion and species richness, we find that species richness (Figs. 1 and 6a) and probability of invasion (Figs. 2 and 6b) are independently modulated by the distribution of interaction types. Our results thus renew focus on the balance of interaction types as a unifying theme for explaining invasibility of ecological networks [56].

**Figure 6:**
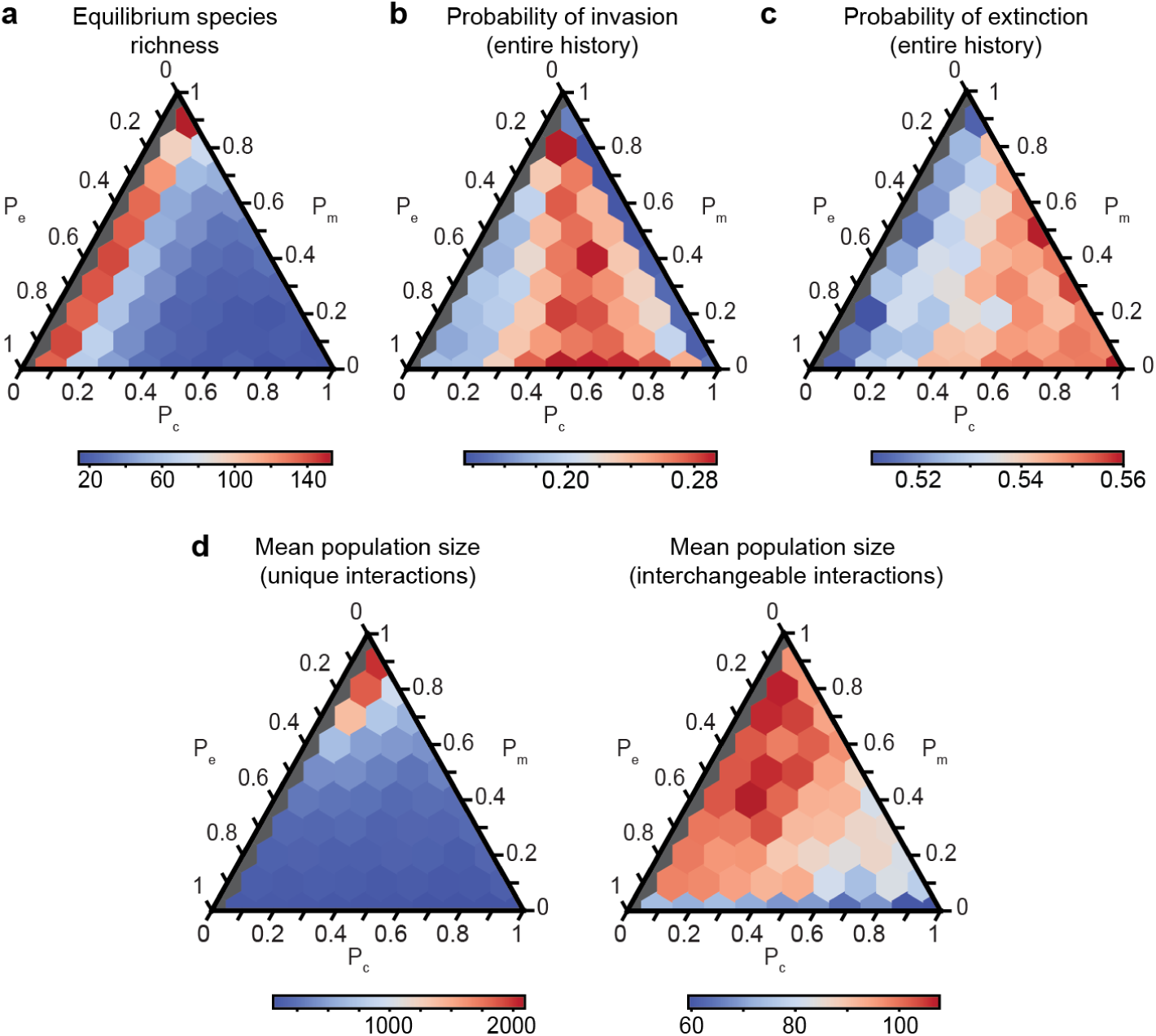
The interchangeable interactions model shows some notable differences compared to the unique interactions model. Panels A through C are for the IIM with *C* = 0.5 and *h* = 100. Gray communities have no competition and as with the UIM, are not considered since they are biologically unrealistic. (a) Steady state species richness of different communities with varying proportions of competition, mutualism, and exploitation. (b) External stability throughout the entire community history. (c) For the IIM with *C* = 0.5 and *h* = 100, probability of extinction throughout the entire community history. These panels show that steady state diversity, external stability, and extinction probabilities mostly depend on the proportion of competitive interactions, with weak effects of the relative balance of exploitative and mutualistic interactions for a given level of competition. (d) Comparison of the mean population sizes of all species in the UIM and IIM with *C* = 0.5 and *h* = 100. The absence of a mutualism effect for external stability in particular is caused by the fact that in the interchangeable interactions model, there is much less variation in population sizes with the proportion of mutualism.

The role of species interactions in community assembly has long been a central concern of ecology [49, 50]. It has also recently become a focus of studies on community stability [10, 13, 14, 18]. Our results suggest that the balance of species interactions are a fundamental driver of community dynamics. Numerous studies have examined the relationships between community stability, invasability, diversity, and saturation, but we argue that species interactions operate at a more elementary level and are the determinants of such other higher-order properties of communities. In particular, mutualisms may play a more important role than suggested by previous studies that assume linearity of its functional response. We show that considering sequentially assembled ecological communities with biologically realistic nonlinear functional responses allows a resolution of the complexity-stability debate and brings into focus other important properties of communities.

## Methods

We construct a model of community assembly to study how communities of desired properties are formed over time. The species richness of a community changes over time and is denoted *S*. The community is initialized with *S*_0_ species and grows in size as successful invasions by new species occur. One of the key features of our model is the ability to consider any combinations and proportions of any interaction types in the construction of the interaction matrix *A* of the community, similar to Coyte *et al.* [14]. We likewise limit our model to mutualism, competition, and exploitation for simplicity, though our model can easily be generalized to include commensalism and ammensalism. As model parameters, we would like to impose a desired connectivity of *A*, denoted *C*, and desired proportions of mutualistic, competitive, and exploitative interactions, denoted by *P_m_*, *P_c_*, and *P_e_*, respectively. The normalization constraint is simply *P_m_* + *P_c_* + *P_e_* = 1.

For the UIM, the population dynamics of species *i* is given by

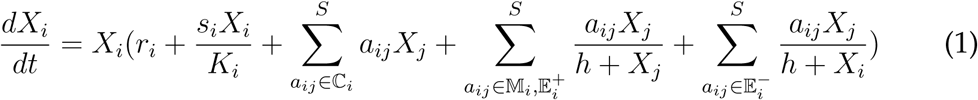

and for the IIM, the population dynamics of species *i* is given by

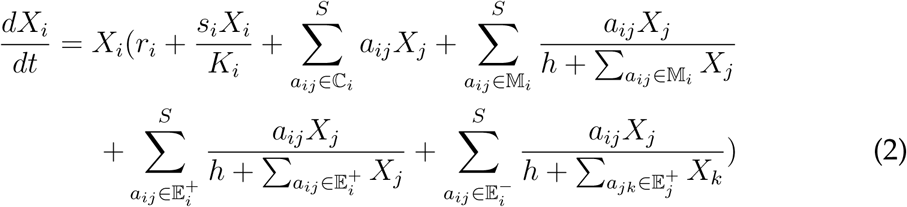

where *X_i_* is the population of species *i*, *r_i_* is its intrinsic growth rate, *a_ij_* is the interaction coefficient between species *i* and *j*, *h* is the half-saturation constant of the Type II functional response, *s_i_* is the density-dependent self-regulation term (and is negative), and *K_i_* is carrying capacity. C*_i_* is the set of interactions between species *i* and its competitors. Likewise, 𝕄*_i_* is the set of interactions between species *i* and its mutualistic partners. 𝔼_i_^+^ refers to the set of interactions between species *i* and the species that it exploits, while 𝔼*^−^_i_* refers to the set of interactions in which species *i* is getting exploited. In equation 1, the separation of the summation over these sets is simply the standard implementation of the predator-prey Type II functional response, where interaction effects for both predator and prey saturate with increasing population of prey. In both the UIM and IIM, we can account for any variation in competition (both *a_ij_* and *a_ji_* are negative), exploitation (*a_ij_* and *a_ji_* are opposite signs), and mutualism (both *a_ij_* and *a_ji_* are positive) throughout the community.

The UIM and IIM are implemented identically except for the population dynamics given above. Each model begins with *S*_0_ species whose population dynamics are governed by equation 1 or 2. Their initial populations are set equal to a constant *x*_0_ multiplied by a random variable drawn from a uniform distribution on [0, 1]. To focus on the effects of interaction types, we set the intrinsic growth rates, density-dependent self-regulation terms, and carrying capacities of all species to be equal, so *r_i_* = *r*, *s_i_* = *s <* 0, and *K_i_* = *K* for all *i*. We start with an *S*_0_ by *S*_0_ interaction matrix *A*_0_, which stores the interaction coefficients of equation 1 or 2. The diagonal values of *A*_0_ are set to *s*. For each pair of species, i.e., each (*i, j*) combination where *i* ≠ *j*, the probability that an interaction exists is given by *C*; i.e., with probability 1 *− C*, we set the interaction coefficients *a_ij_* = *a_ji_* = 0. For each species pair *i* and *j* that does interact, the interaction is mutualistic with probability *P_m_*, in which case the interaction coefficients *a_ij_* and *a_ji_* are both drawn from a half-normal distribution *N* (0*, σ*) where *σ* is the scale parameter. Likewise, with probability *P_c_*, *i* and *j* compete with each other, in which case both *a_ij_* and *a_ji_* are drawn from a negative half-normal distribution *−N* (0*, σ*/*K*), where the scale parameter is scaled by the carrying capacity. This is such that the average effect of competition on population dynamics is comparable to that of mutualism. Finally, with probability *P_e_* = 1 *− P_m_ − P_c_*, *i* and *j* engage in an exploitative interaction. In this case, we assign one of *i* or *j* randomly as the exploited species, while the other becomes the exploiter. If species *i* is the exploiter, *a_ij_* is drawn from the half-normal distribution *N* (0*, σ*), and *a_ji_* from *−N* (0*, σ*), and vice versa if species *j* is the exploiter.

The construction of *A*_0_ follows the approach developed by Coyte *et al.* [14]. We then add into the model community assembly and a different manner of simulating population dynamics. After *A*_0_ is created, we simulate the population dynamics of all *S*_0_ species until the populations come to an equilibrium. Integration of the Lotka-Volterra dynamics occurs with adaptive step sizes using the “Dopri5” integrator [57]. Equilibrium of populations in our model can be achieved in two ways. First, we impose a population change threshold *δ*, so that if in a single time step every single population fluctuates by less than *δ*, then the populations are determined to have equilibrated. Second, for simulation purposes we impose a time limit *t_l_* so that if the populations still have not equilibrated after *t_l_* time steps, we use the current population sizes as the equilibrium values. We choose large enough *t_l_* so this does not occur unless populations have entered a stable limit cycle and are oscillating. As species populations change in size, some species may go extinct. We define an extinction threshold *ɛ* so that any species whose population *X_i_* dips under *ɛ* is considered extinct and removed from the community at the next equilibrium.

After the populations have equilibrated, we add a new species *z* into the community, so that *A* increases in size from *S*_0_ by *S*_0_ to *S*_0_ + 1 by *S*_0_ + 1 (in the case that no species have gone extinct in the first iteration of the model). Entries in the new row and column that are added to the interaction matrix are drawn in the exact same manner as described above for the initialization of *A*_0_, so the desired properties *C*, *P_m_*, *P_c_*, and *P_e_* are preserved as the community changes in size. Importantly, not every species introduced randomly in this manner will be able to invade the community. In order for successful invasion to occur, the growth rate of *z* must be positive when a small but nonzero number of species *z* individuals are suddenly added into the community. If the invasion condition is not met and a failed invasion occurs, we replace the failed invader with a new species whose interaction coefficients are drawn anew in the same aforementioned manner. We repeat this process until a new species is able to invade the community. We track the number of failed invasions. For computational tractability, we impose a failed invasion threshold *β* so if over *β* new species fail to invade a community at any given equilibrium, the community is deemed externally stable and the simulation ends. This scenario is not reached in the communities we simulate.

On the other hand if the invasion condition is met, the community grows by one species. The new species is initialized with a small population size equal to a constant *x_z_* multiplied by a random variable drawn from a uniform distribution on [0, 1]. We then simulate the population dynamics until the next equilibrium, at which point we introduce another new species, and so on. In this manner, at each equilibrium a new species is added into the community. The species richness and probabilities of invasion and extinction are tracked. We impose a limit *n* on the species richness of a community for computational tractability so if *S* exceeds *n*, the simulation ends (this does not occur in our simulations). As mentioned in the main text, communities exhibit a convergence of species richness. The presence of a steady state of species diversity is assessed using an augmented Dickey-Fuller test, a unit root test whose null hypothesis is that the species richness time series has a unit root (i.e., is non-stationary and has time-dependent structure) and whose alternative hypothesis is that species richness does not have a unit root and is stationary over time. We analyze the species richness of the most recent *w*_1_ equilibria of the community and reject the null hypothesis when the p-value is below some threshold *ρ*. Rejection of the null hypothesis indicates that species richness has entered a steady state. If a community enters the steady state, the simulation runs for *w*_2_ more equilibria before it is terminated. Simulations with communities that never enter the steady state can terminate in one of three ways: by reaching the limit on species richness (*n*), the limit on the number of equilibria (*κ*), or the time limit for the overall simulation (*t_t_*), though in our simulations these limits are never reached.

For each combination of *C* and *h*, we simulated communities by varying the proportions of competition, mutualism, and exploitation from 0 to 1 in intervals of 0.1, subject to the condition that their sum is equal to 1; there are 66 such combinations of interaction types, and we simulate each combination with five replicates to get 330 communities. Since there are 9 combinations of *C* and *h*, there are a total of 2970 communities. We discard communities with *P_c_* = 0 during our analysis, since communities at this edge case do not saturate in species richness. Since real communities are expected to have at least some competition, this does not affect our conclusions. Supplementary Table 1 summarizes the parameters we used in our simulations. We choose *s* and *σ* such that the magnitude of the intraspecific interaction is generally larger than the magnitude of interspecific interaction. This biologically realistic assumption easily arises from the fact that members of the same species compete more strongly for the same set of resources and space than do members of different species. In the main text, we use *C* = 0.5 and *h* = 100, while in the Supplementary Information we extend our analysis to other values of *C* and *h*. We perform the full analysis for both the UIM and IIM.

All data shown in figures in the main text are averages over five replicates, except for Figs. 1a and 4a, which show data from representative communities, and Fig. 5. To study selection (Fig. 5), we simulate communities until species diversity converges to a steady state. We take advantage of the fact that the communities appear to exhibit ergodicity (as suggested by Fig. 1 and Supplementary Fig. 2). We continue to randomly create new species and allow the community to come to 100 *S_s_* additional equilibria, where *S_s_* is the steady state species diversity. At each equilibria, species are introduced until successful invasion occurs. We sample the community 100 times at an interval of every *S_s_* equilibria and calculate the average distribution of interaction types (Fig. 5a). After the steady state is reached, we save the interaction types of every tenth successful invader and every tenth species that goes extinct. The average interaction proportions are calculated and plotted (Fig. 5b,c,d).

## Code availability

The model of community assembly was implemented in Python. The simulation code is available at https://github.com/erolakcay/CommunityAssembly.

## Data availability

Data sharing not applicable to this article as no datasets were generated or analysed during the current study. All simulation results can be reproduced using the code provided.

## Acknowledgments

We would like to thank A. Tilman, J. Van Cleve, L. Stone, and G. Barabás for helpful comments regarding the manuscript. J.Q. was funded by the Roy and Diana Vagelos Scholars Program in the Molecular Life Sciences at the University of Pennsylvania.

## Author contributions

Both authors designed the study. J.Q. constructed the model and both authors provided analysis. Both authors contributed substantially to writing the manuscript. Both authors gave final approval for publication.

## Supplementary Information

The results in the main text were derived from the model with *C* = 0.5 and *h* = 100 (for reference, the carrying capacity *K* = 100 for all simulations). We repeat the analysis with a lower bound *C* = 0.1 and upper bound *C* = 0.9, and lower and upper bounds of *h* = *K*/5 = 20 and *h* = 5*K* = 500, respectively, for both the UIM and IIM. We find that the results are qualitatively robust across all values of *C* and *h* and both the UIM and IIM. Most of the supplementary figures correspond to the figures from the main text, extended to show results for all nine combinations of *C* and *h* for both models.

Supplementary Fig. 1 shows that there is a large difference in population sizes between the UIM and IIM: normalization of interchangeable interactions decreases the population sizes drastically compared to unique interactions.

Supplementary Figs. 2 through 18 show data for the UIM. Supplementary Fig. 2 shows that community saturation occurs for all levels of community connectivity and all half-saturation constants. Supplementary Fig. 3 shows how the steady state species richness is affected by the balance of interaction types. The lower the connectivity, the higher the steady state species richness, but the qualitative pattern of convergence to the steady state is the same. There is not a strong dependence of species richness on *h*. The manner in which the various interaction types affect community saturation follow the patterns described in the main text for all *C* and *h*.

Supplementary Fig. 4 shows representative oscillations during the rare cases when the dynamics fail to converge to an equilibrium. In principle, a new species invading a stable limit cycle is not different than invading the equilibrium. Supplementary Fig. 5 shows that the probability that oscillations occur is very low for all communities. However, when *h* and *P_m_* are both low there can be more oscillations. Only a very small number of communities show probabilities of oscillations that are larger than 0.01, confirming that focusing on external stability rather than internal stability is critical.

Supplementary Figs. 6, 7, and 8 show how external stability is affected by combinations of interaction types across combinations of *C* and *h*. The pattern described in the main text is qualitatively robust across *C* and *h*. However, there is a slight change in the pattern for how interaction types affect external stability as *h* changes (but not when *C* changes). *C* does affect how the probability of invasion changes as the community moves from the transient state into the steady state: when *C* is low, a small subset of communities (low *P_c_*, high *P_m_*) may show a large decrease in invasion of probability (Supplementary Fig. 8), whereas probability of invasion usually does not change upon arrival at the steady state. Supplementary Figs. 9, 10, and 11 show how probability of extinction is governed by the balance of interaction types across combinations of *C* and *h*. Supplementary Fig. 9 shows how throughout the entire community history, the pattern of the effect of interaction types slightly changes with *h*. When *h* is small, the most extinction occurs with low competition, low mutualism, and high exploitation, whereas when *h* is high the most extinction occurs when competition is high and mutualism and exploitation are low. After arrival at the steady state, the probability of extinction increases regardless of *C* or *h* (Supplementary Fig. 10 and 11). When *h* is high, there is a very obvious pattern for how the balance of interaction types modulates the increase in extinction.

Supplementary Figs. 12, 13, and 14 show how species persistence is determined by the proportion of interaction types. For all *C* and *h*, the patterns are the same as described in the main text. Low connectivity allows for longer persistence and less species turnover, but the effect of interaction types on persistence are conserved across all levels of connectivity. Higher *h* also allows for longer species persistence especially when *P_m_* is lower.

Supplementary Figs. 15, 16, 17, and 18 show how sequential selection determines community composition and modulates the distribution of interaction types. For all *C* and *h*, selection increases the amount of mutualism in the community. Connectivity does not exert an influence on the distribution of realized interaction types, but interestingly the half-saturation constant does. For *h* = 500, selection clearly acts to increase mutualism, decrease competition, and slightly increase exploitation, with a more dramatic increase in exploitation at lower levels of mutualism. Species that successfully invade each community show the same bias in their interaction types, implying that selection acts at the level of invasion. In general, species that went extinct showed the same biases in interaction proportions as successful invaders. There is no clear difference in the interaction types between species that invade and those that go extinct, suggesting that selection for some species to persist longer than others relies on the topology of the interspecies links (i.e. community architecture) rather than the types of those interactions. The *h* = 100 case is discussed in detail in the main text. On the other hand, the *h* = 20 case shows a somewhat different pattern. For all values of *h*, selection acts to increase mutualism, but when *h* = 20 it also slightly decreases exploitation especially when there is high exploitation. Moreover, selection increases competition when it is low, and decreases competition when it is high. This is different than the *h* = 500 case when competition is always decreased. The increase in competition is also present for more combinations of interaction types than compared to the *h* = 100 case. When *h* = 20, the bias of interaction types between species that invade and those that go extinct are different: when competition is low and exploitation is high, species that go extinct tend to have less competition than species that invade the community.

Supplementary Figs. 19 to 33 show the corresponding data for the IIM.

**Supplementary Table 1:** Summary of model parameters that were held constant across all simulations in the main text.

| Parameter | Description |
| --- | --- |
| $r = 1$ | intrinsic growth rate of all species |
| $K = 100$ | carrying capacity of each species |
| $h = 20$ | half-saturation constant for Type II functional response |
| $s = -1$ | self-regulation coefficient (diagonals of the interaction matrix) |
| $S_0 = 10$ | number of species in initial community |
| $x_0 = 10$ | maximum initial population size of original species |
| $x_z = 10$ | maximum initial population size of introduced species |
| $n = 750$ | limit on species richness |
| $\kappa = 5000$ | limit on number of equilibria |
| $C = 0.5$ | desired connectivity of the ecological network |
| $\sigma = 0.5$ | half-normal distribution scale parameter |
| $\delta = 0.001$ | equilibration criteria for changes in population |
| $t_l = 5000$ | time limit to wait for equilibrium |
| $\epsilon = 0.01$ | extinction threshold for population sizes |
| $\beta = 1000$ | failed invasion threshold |
| $w_1 = 1000$ | number of equilibria analyzed in augmented Dickey-Fuller test |
| $w_2 = 500$ | number of equilibria simulated after steady state is reached |
| $\rho = 0.01$ | $p$-value threshold to reject null hypothesis |
| $t_t = 10^9$ | time limit of simulation |

**Supplementary Figure 1:**
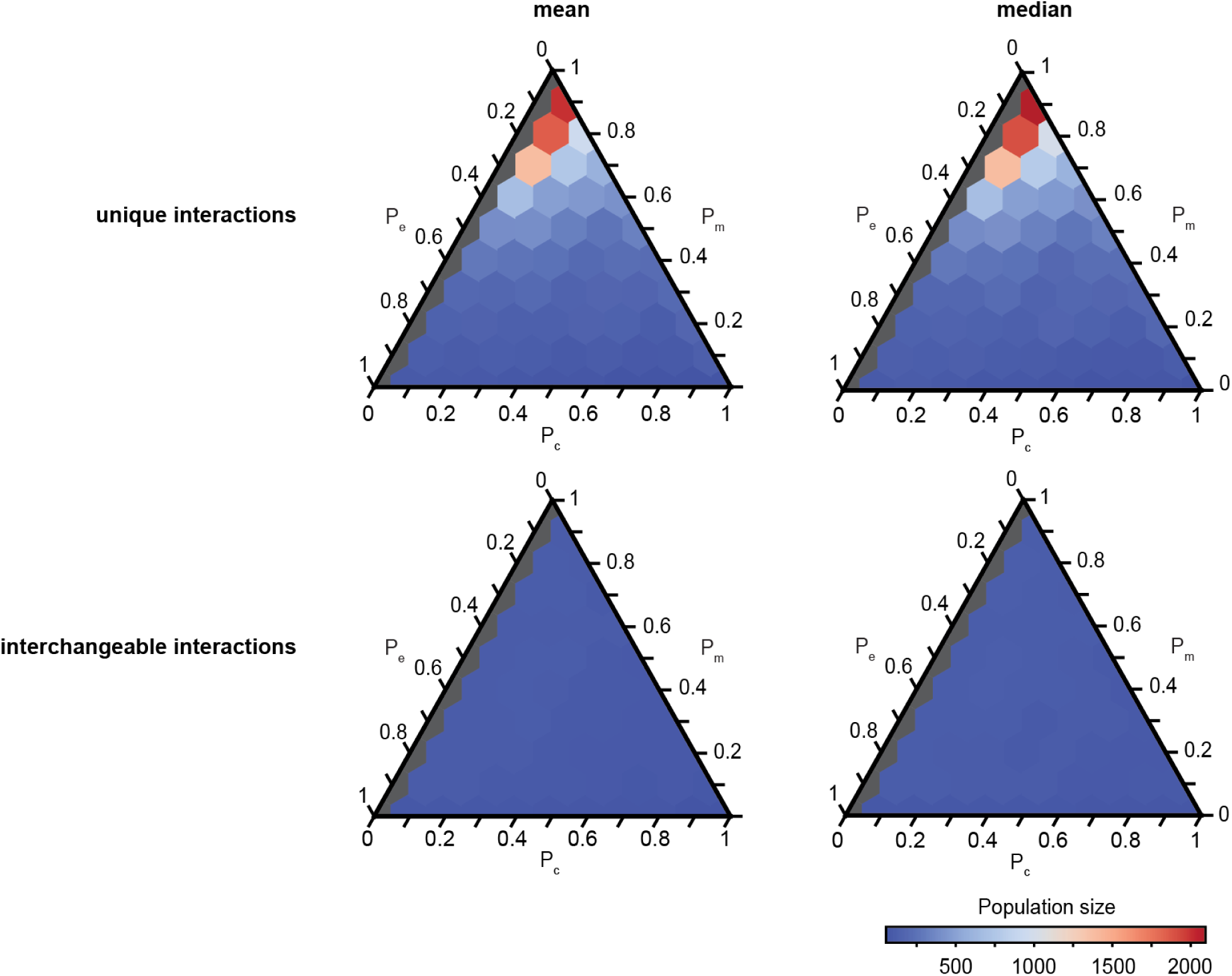
Comparison of the mean and median equilibrium species population sizes for communities of all combinations of interaction types, for both the UIM and IIM (with representative choice *C* = 0.5 and *h* = 100). The colors in all plots correspond to the color bar shown.

**Supplementary Figure 2:**
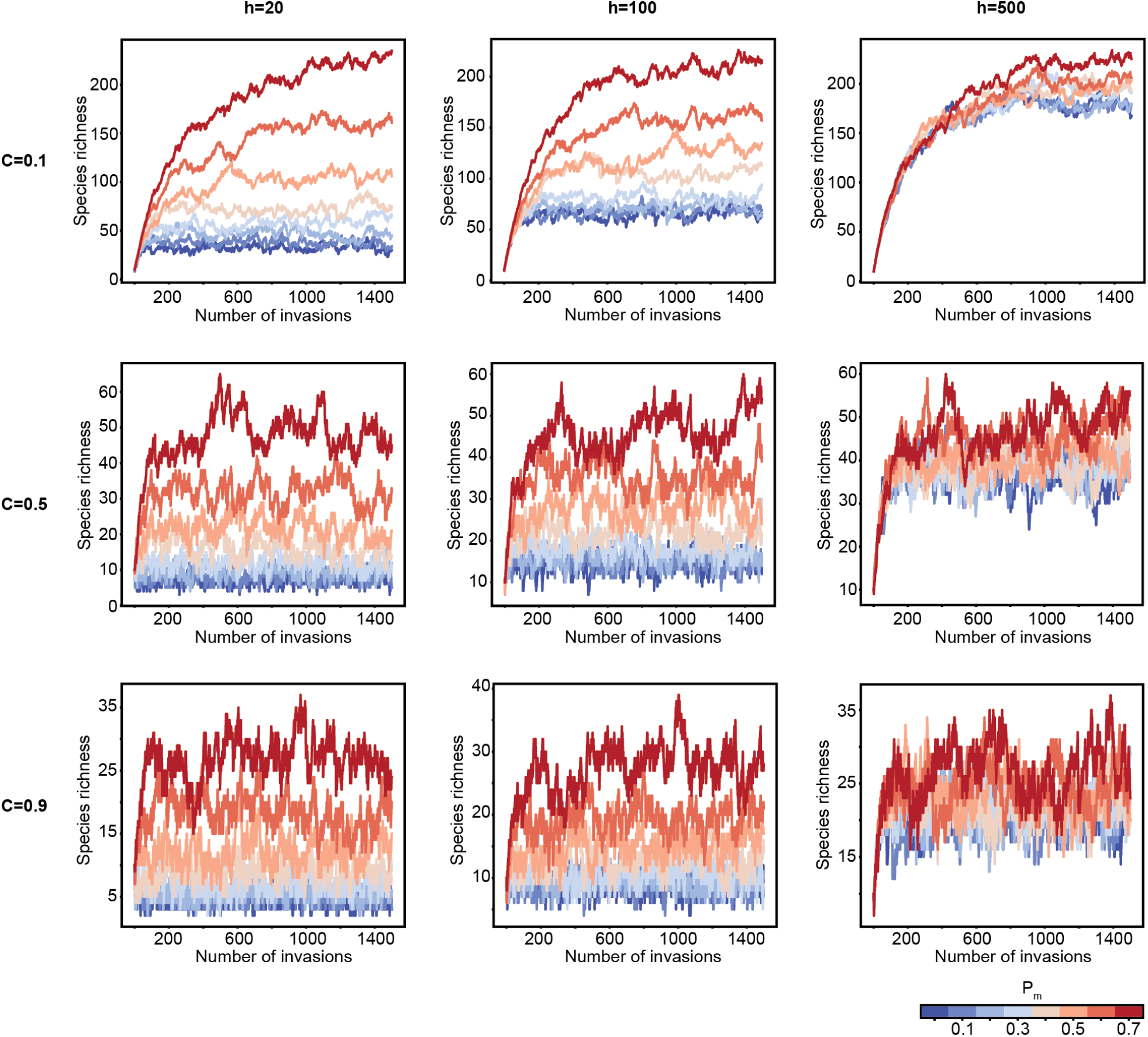
For the UIM, the mixture of interaction types determines the species richness at which communities saturate, for various *C* and *h*. Species richness in representative communities with *P_c_* = 0.3 and varying levels of *P_m_*, represented by the color. The colors in all plots correspond to the color bar shown. Each line corresponds to a community over 1500 species invasions that occur at equilibria. The same trend is observed for all *P_c_ >* 0.

**Supplementary Figure 3:**
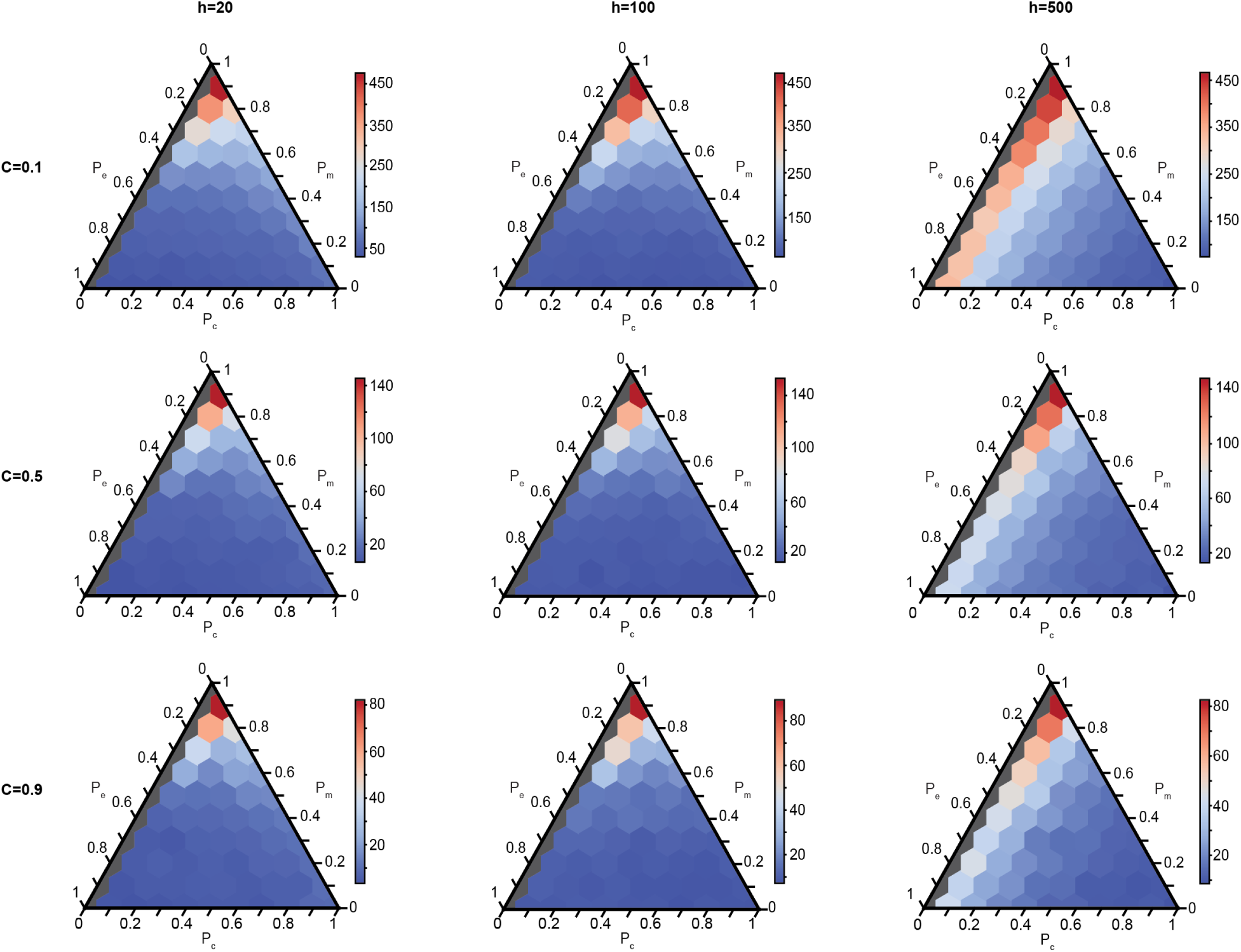
For the UIM, the steady state species richness of different communities with varying proportions of competition, mutualism, and exploitation, and varying *C* and *h*. The color represents the species richness, and each plot has its own color scale. Gray communities have no competition and are not considered since they are biologically unrealistic.

**Supplementary Figure 4:**
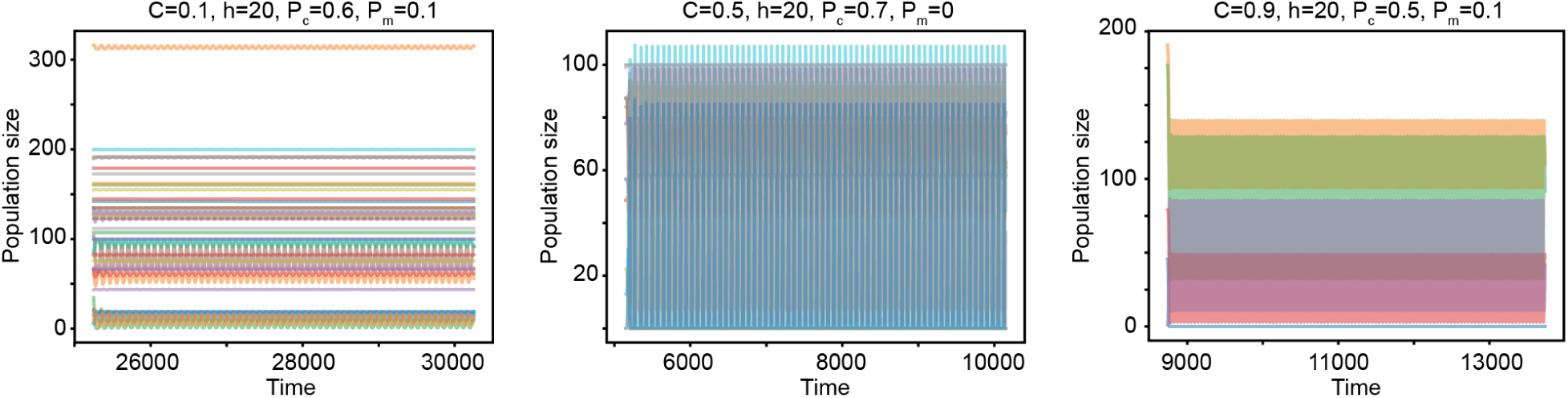
For the UIM, representative oscillations that occur when the dynamics fail to converge to an equilibrium. Each line corresponds to one species. When oscillations occur, the simulation introduces a new species that can possibly invade, subject to the invasion condition, once the time limit for equilibration is reached. The time shown in the plots is the integration time of the community history and spans the time limit for equilibration.

**Supplementary Figure 5:**
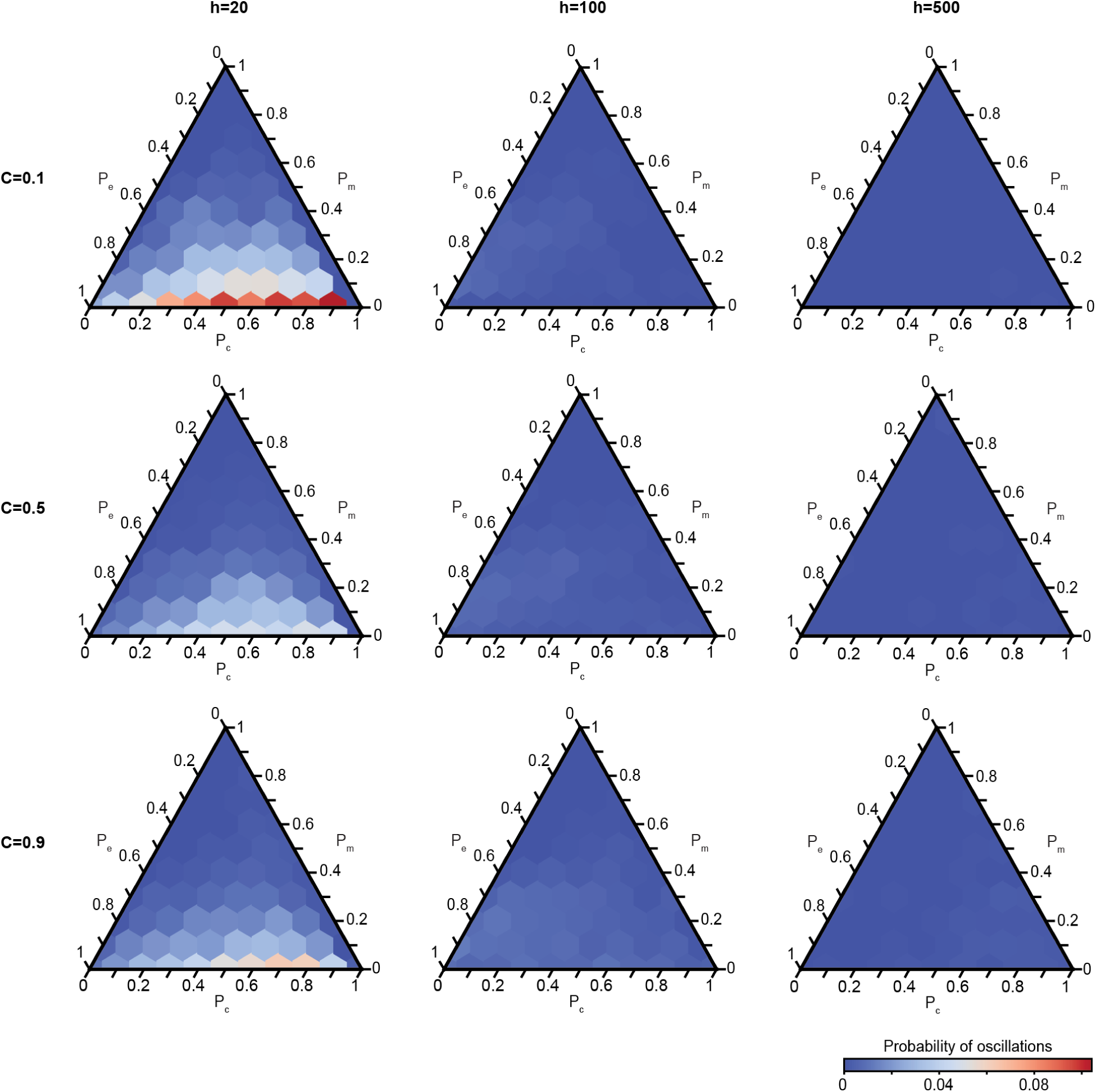
For the UIM, the probability of oscillations throughout the entire community history, for all combinations of interaction types, *C*, and *h*. The color represents the probability and all plots use the same scale, following the color bar shown.

**Supplementary Figure 6:**
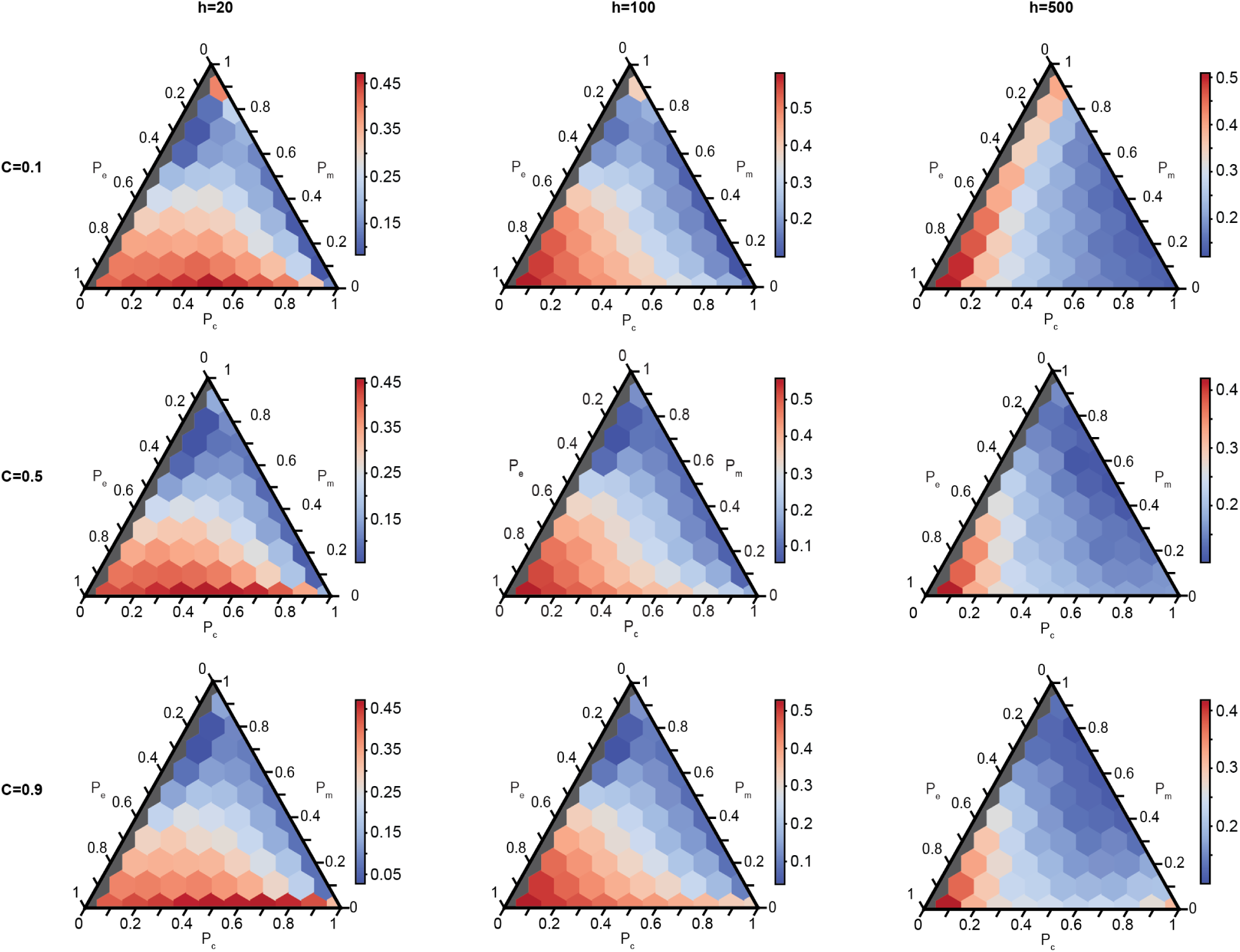
External stability throughout the entire community history for the UIM for varying *C*, *h*, and proportion of interaction types. Colors indicate the probability of invasion and each plot has its own color scale. Gray communities have no competition and are not considered.

**Supplementary Figure 7:**
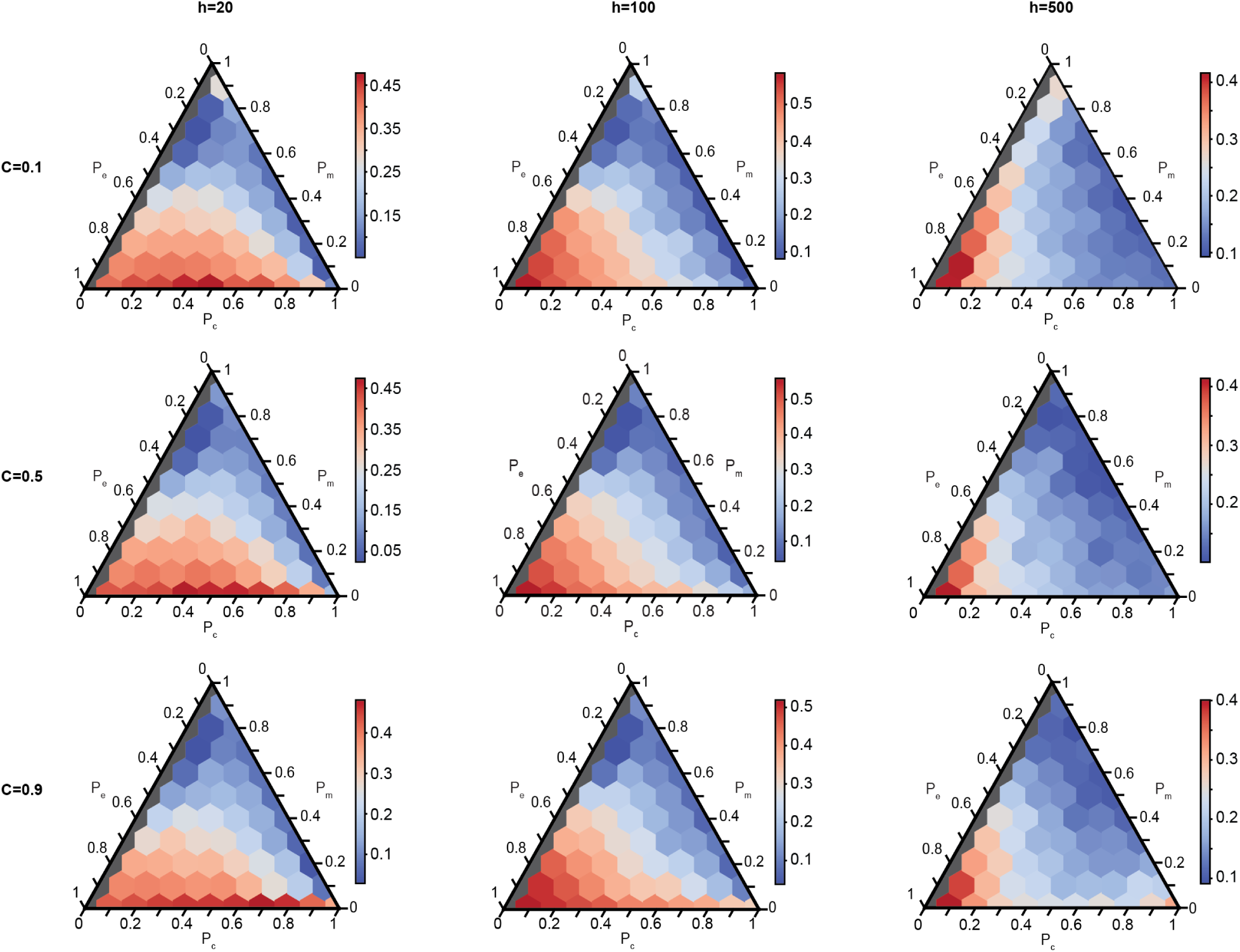
External stability after species richness has converged to the steady state for the UIM with varying *C*, *h*, and proportion of interaction types. Colors indicate the probability of invasion and each plot has its own color scale. Gray communities have no competition and are not considered.

**Supplementary Figure 8:**
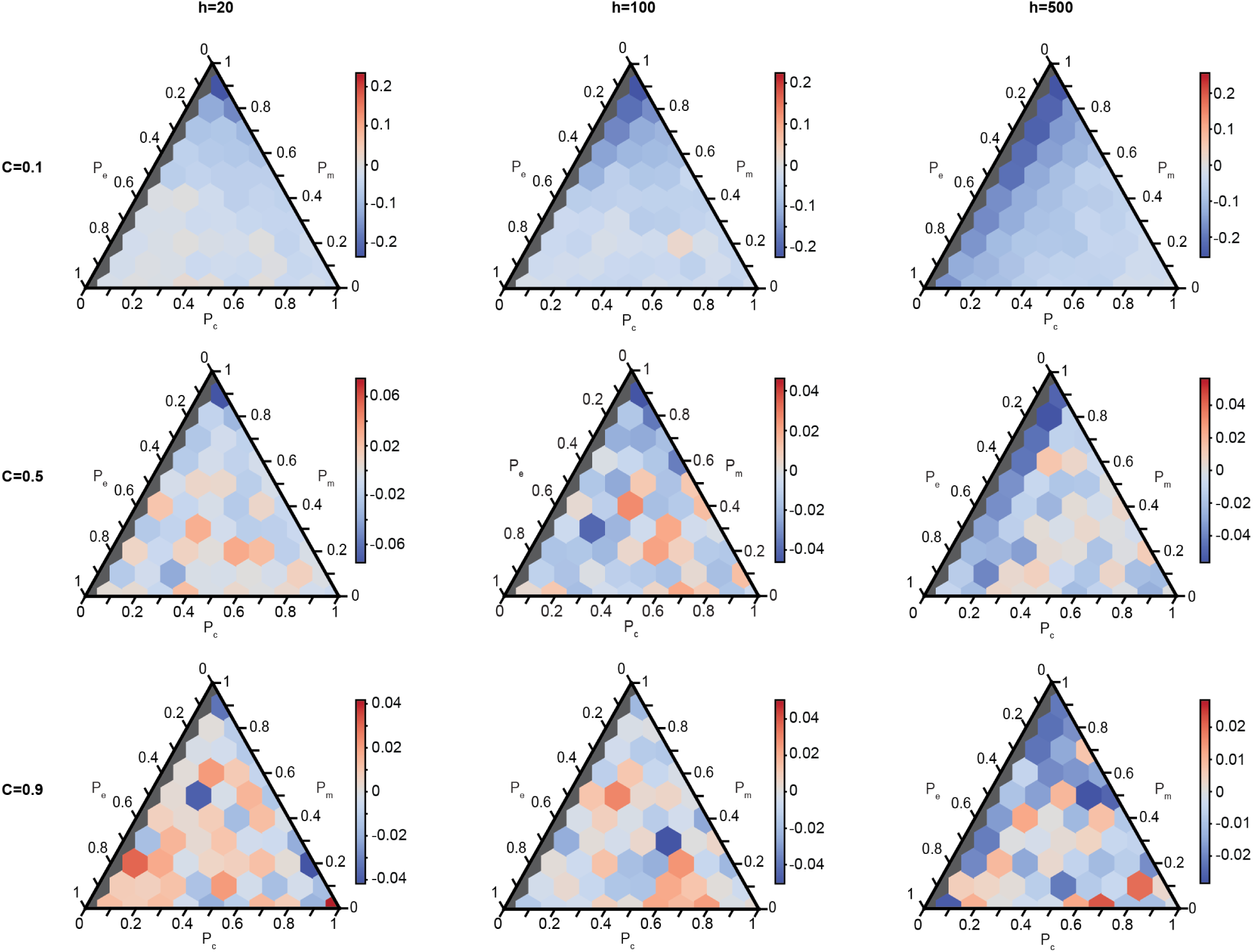
The difference in probability of invasion between the transient state and the steady state for the UIM for varying *C*, *h*, and proportion of interaction types. Colors indicate the change in the probability of invasion and each plot has its own color scale. Gray communities have no competition and are not considered.

**Supplementary Figure 9:**
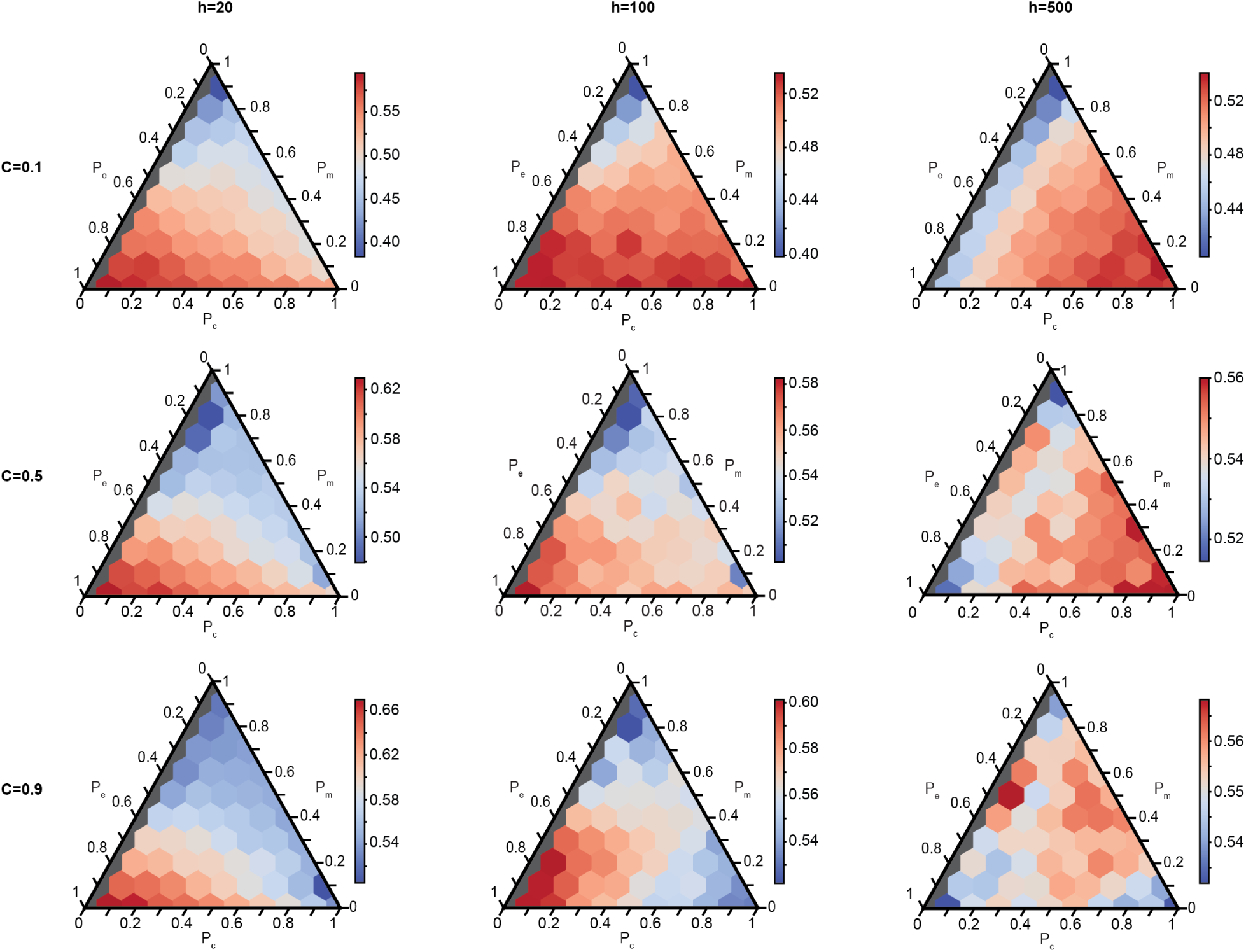
Probability of extinction throughout the entire community history for the UIM for varying *C*, *h*, and proportion of interaction types. Colors indicate the probability of extinction and each plot has its own color scale. Gray communities have no competition and are not considered.

**Supplementary Figure 10:**
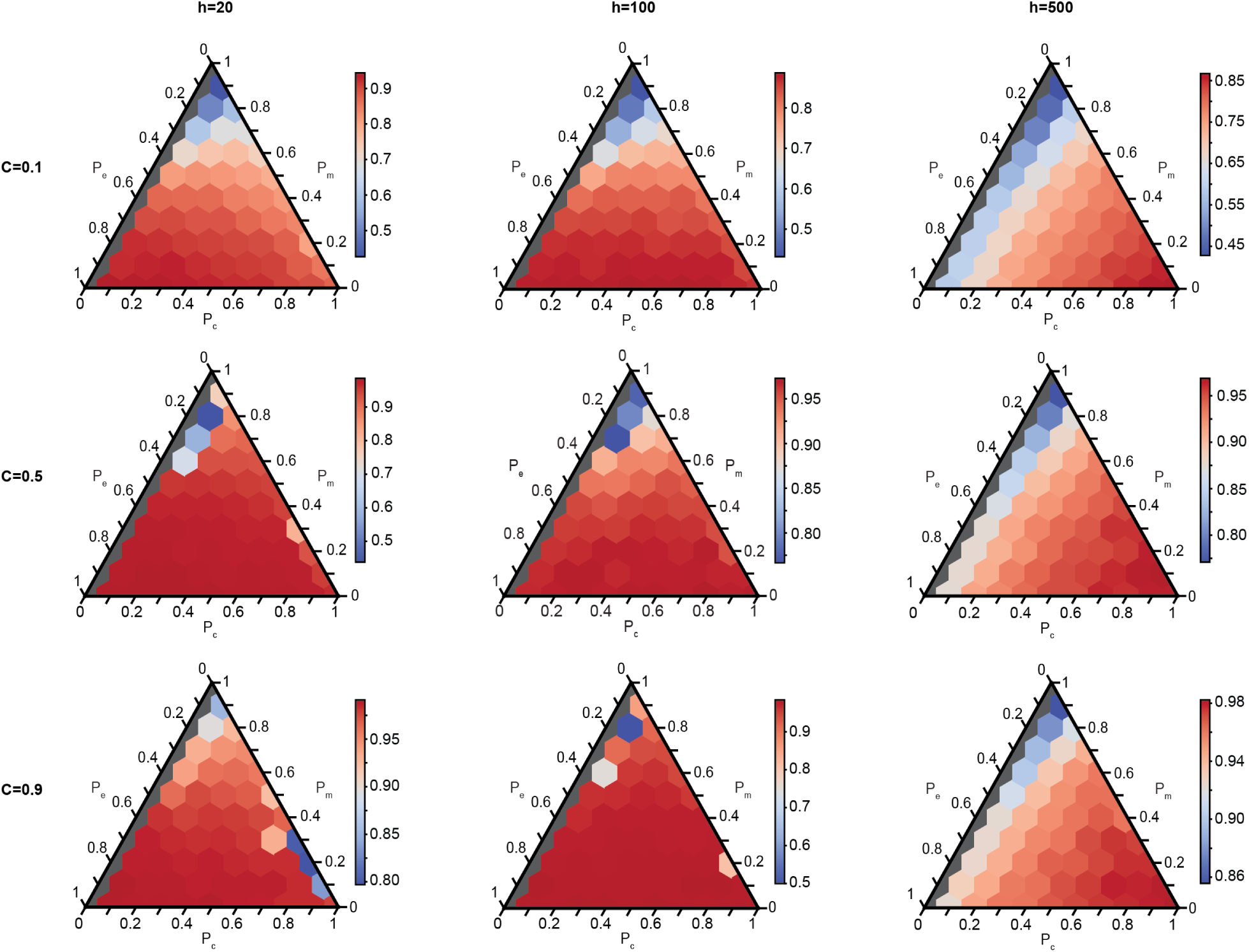
Probability of extinction after species richness has converged to the steady state for the UIM for varying *C*, *h*, and proportion of interaction types. Colors indicate the probability of extinction and each plot has its own color scale. Gray communities have no competition and are not considered.

**Supplementary Figure 11:**
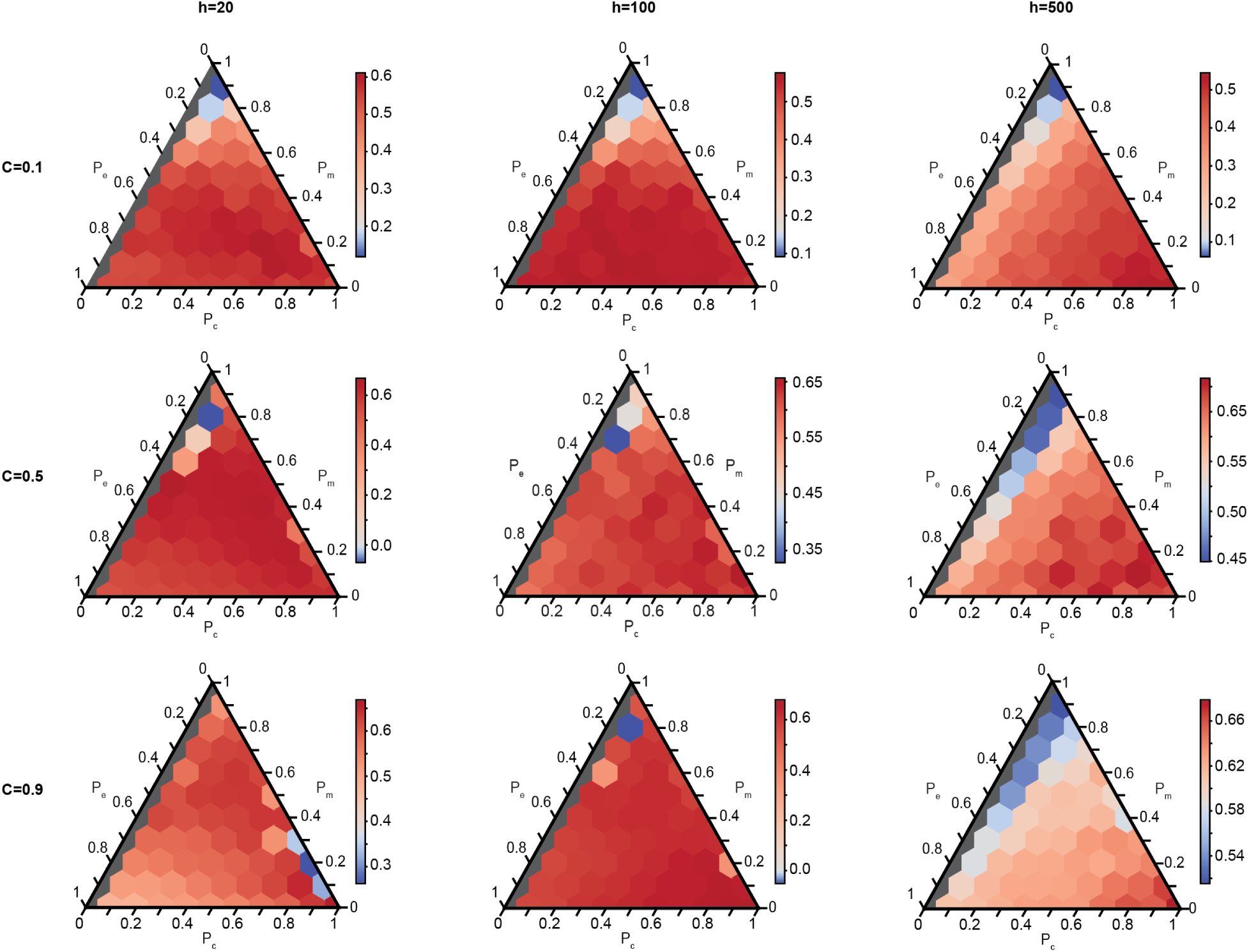
The difference in probability of extinction between the transient state and the steady state for the UIM for varying *C*, *h*, and proportion of interaction types. Colors indicate the change in the probability of extinction and each plot has its own color scale. Gray communities have no competition and are not considered.

**Supplementary Figure 12:**
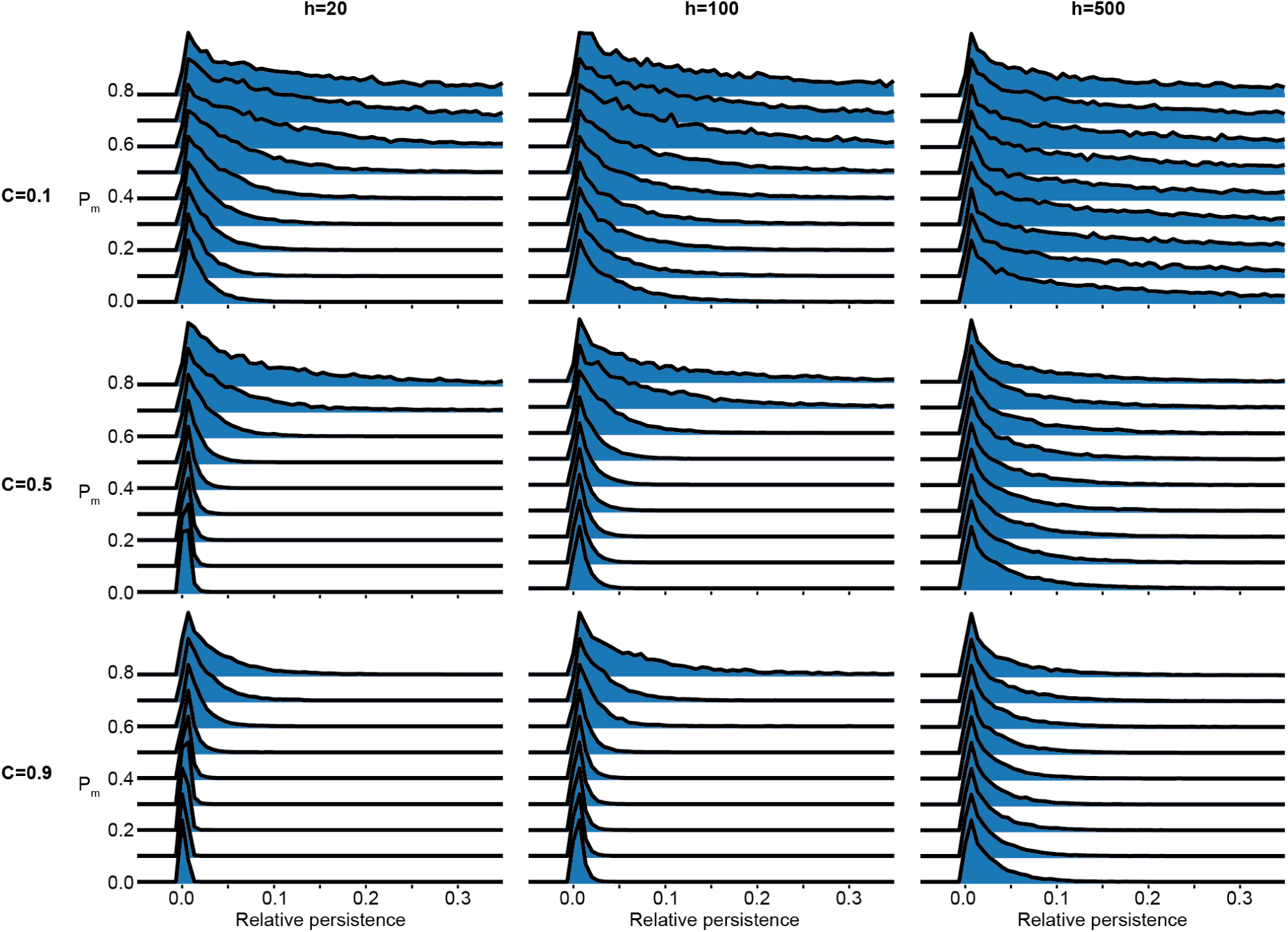
Normalized histograms of the relative persistence of species in communities with *P_c_* = 0.1 and varying *P_m_* for the UIM for various *C* and *h*. This same trend is observed for all *P_c_*.

**Supplementary Figure 13:**
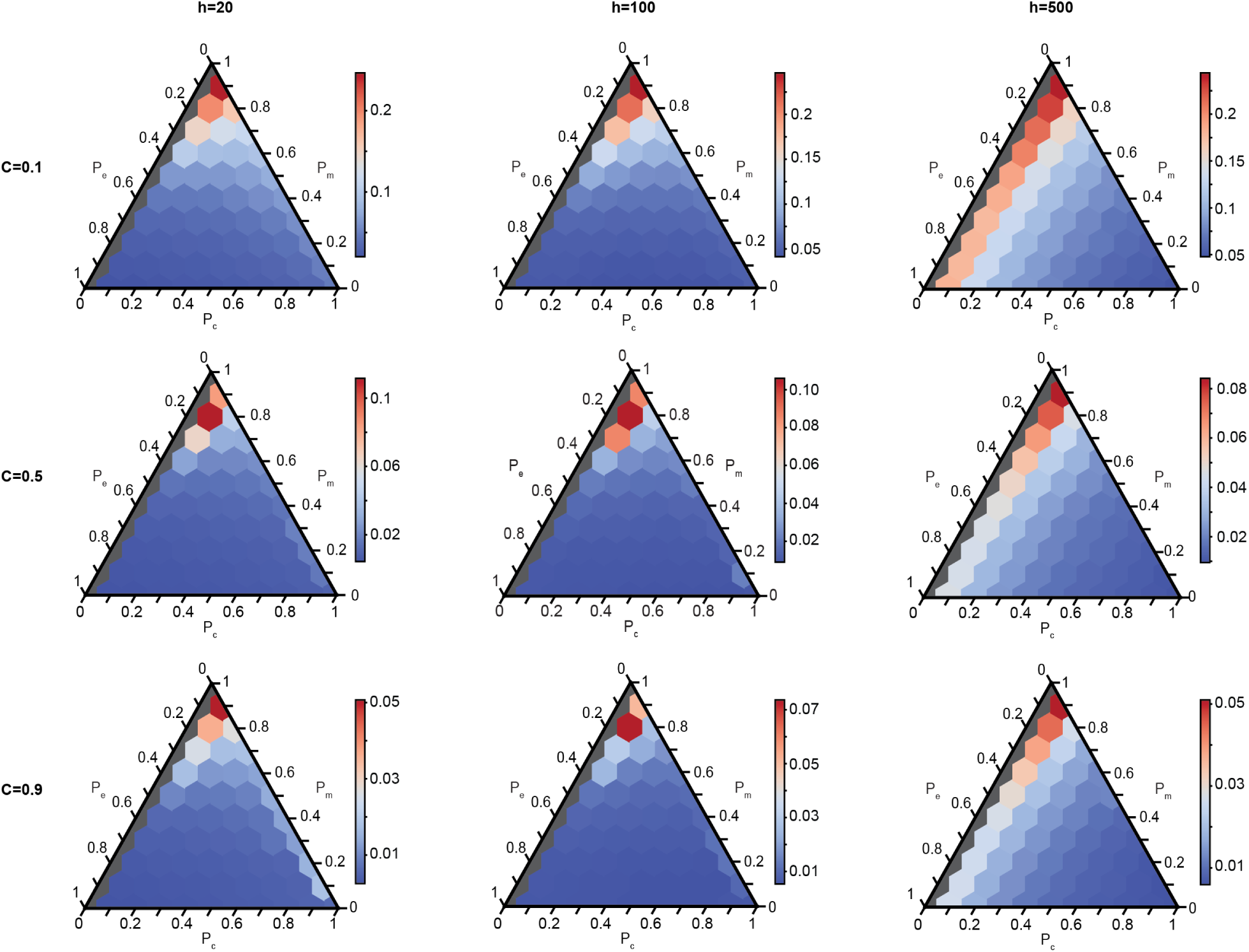
The mean relative persistence of all species that existed within a community for the UIM for varying *C*, *h*, and proportion of interaction types. The color represents the mean relative persistence, and each plot has its own color scale. Gray communities have no competition and are not considered.

**Supplementary Figure 14:**
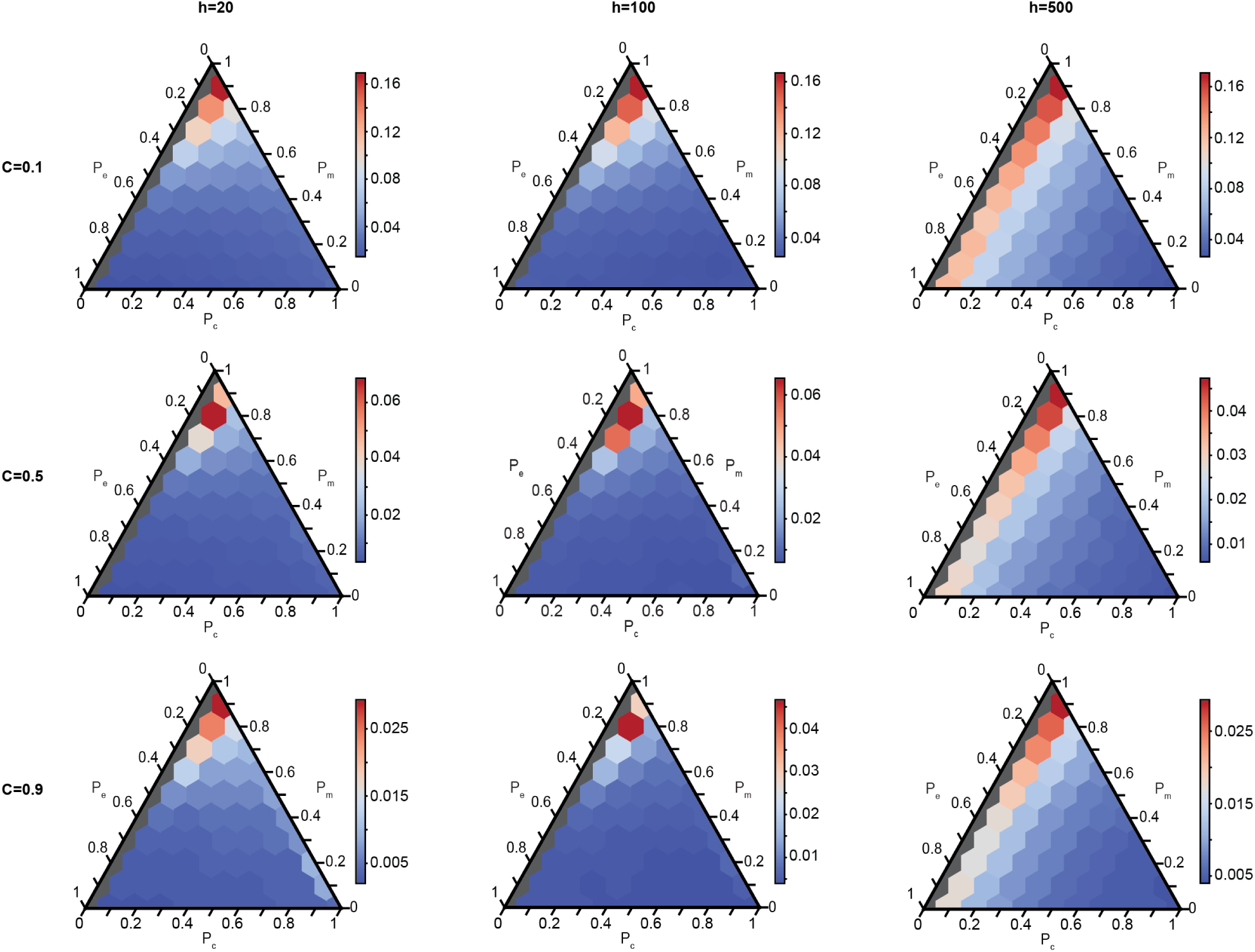
The median relative persistence of all species that existed within a community for the UIM for varying *C*, *h*, and proportion of interaction types. The color represents the median relative persistence, and each plot has its own color scale. Gray communities have no competition and are not considered.

**Supplementary Figure 15:**
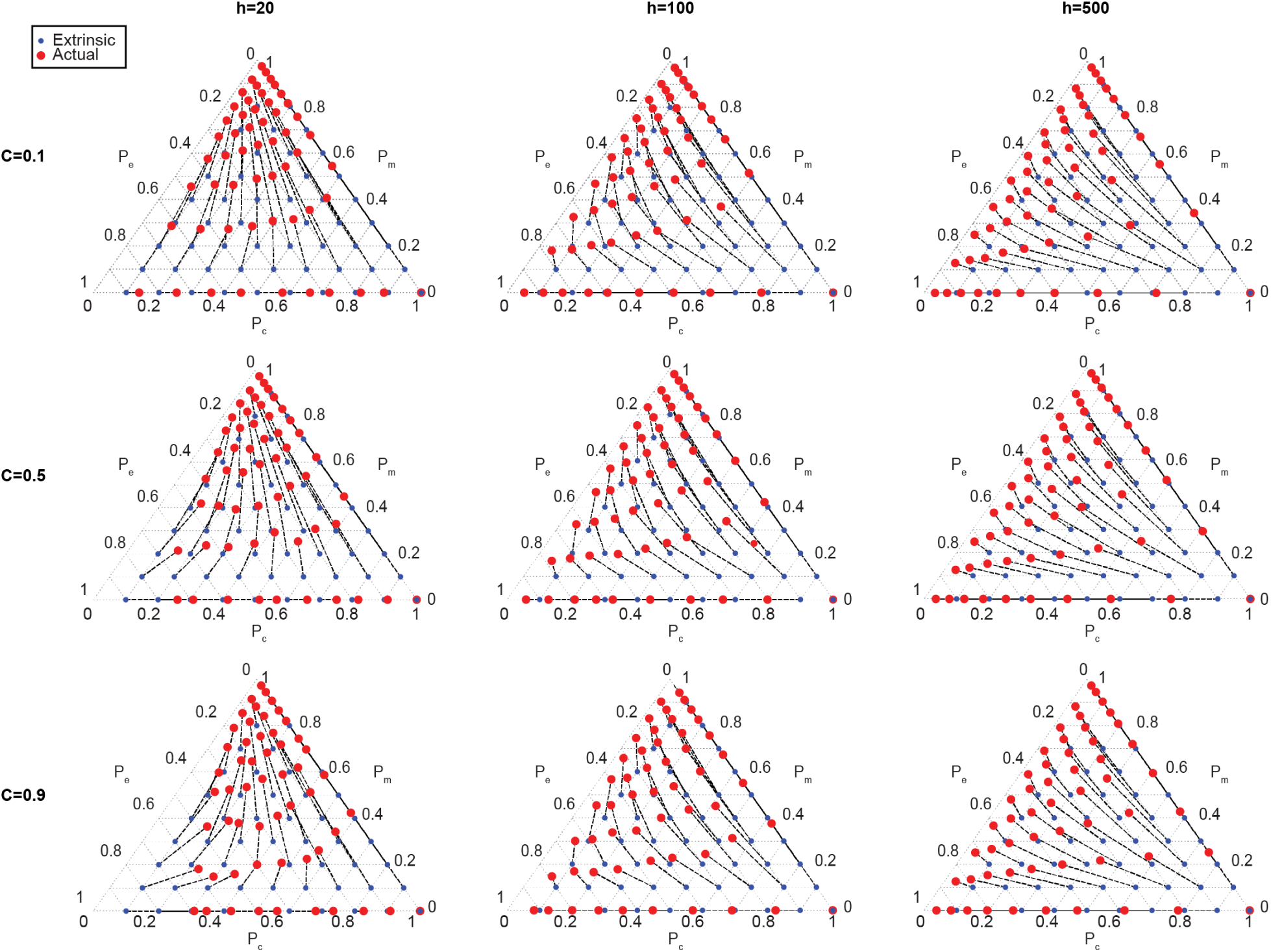
The extrinsic (blue points) and the actual (red circles) interaction proportions of all species within each community for the UIM for varying *C* and *h*. Dotted lines connect points from the same community, and can be interpreted as the direction of selection.

**Supplementary Figure 16:**
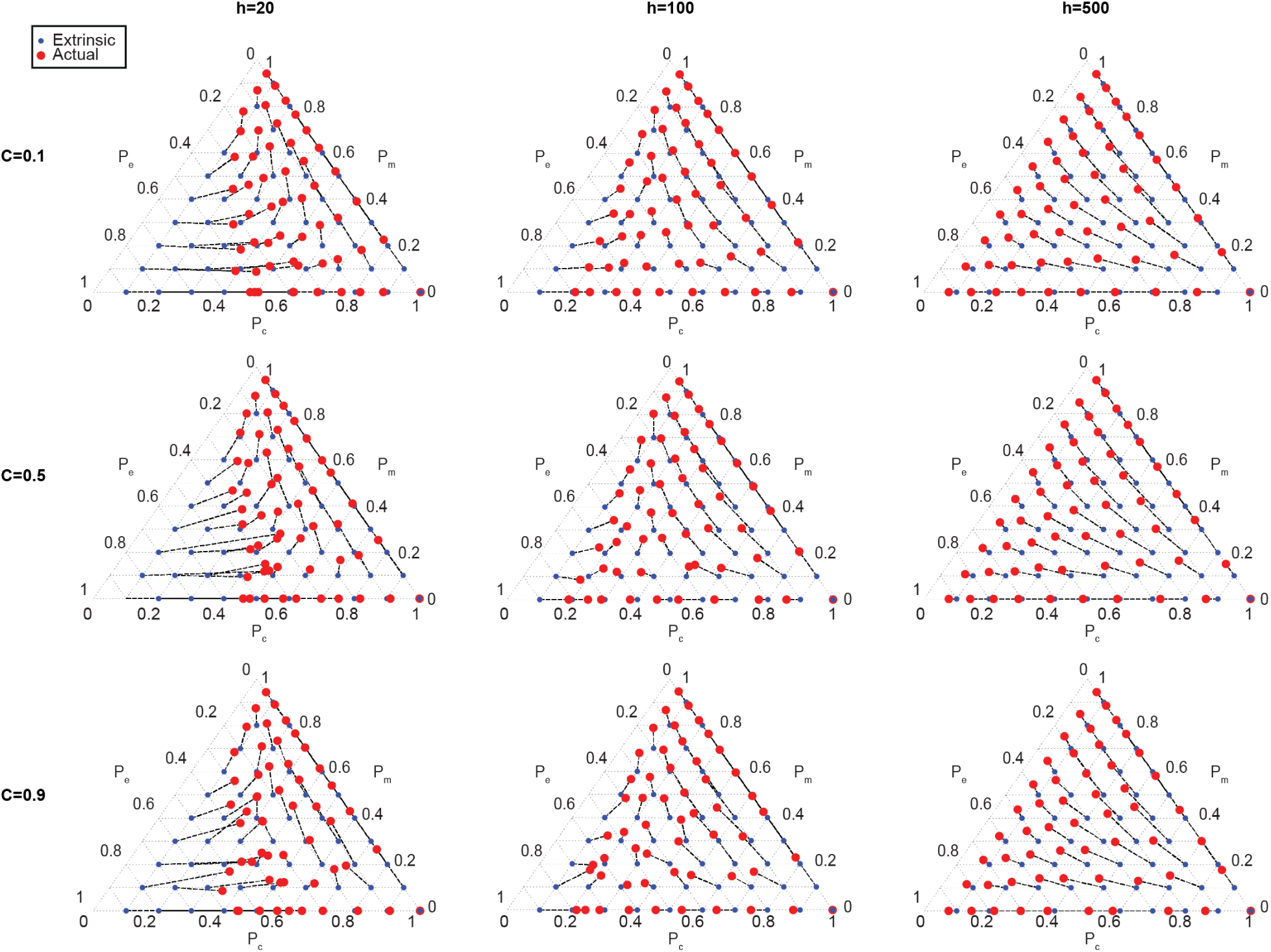
The extrinsic (blue points) and the actual (red circles) interaction proportions of species that successfully invade each community for the UIM for varying *C* and *h*. Dotted lines connect points from the same community, and can be interpreted as the direction of selection.

**Supplementary Figure 17:**
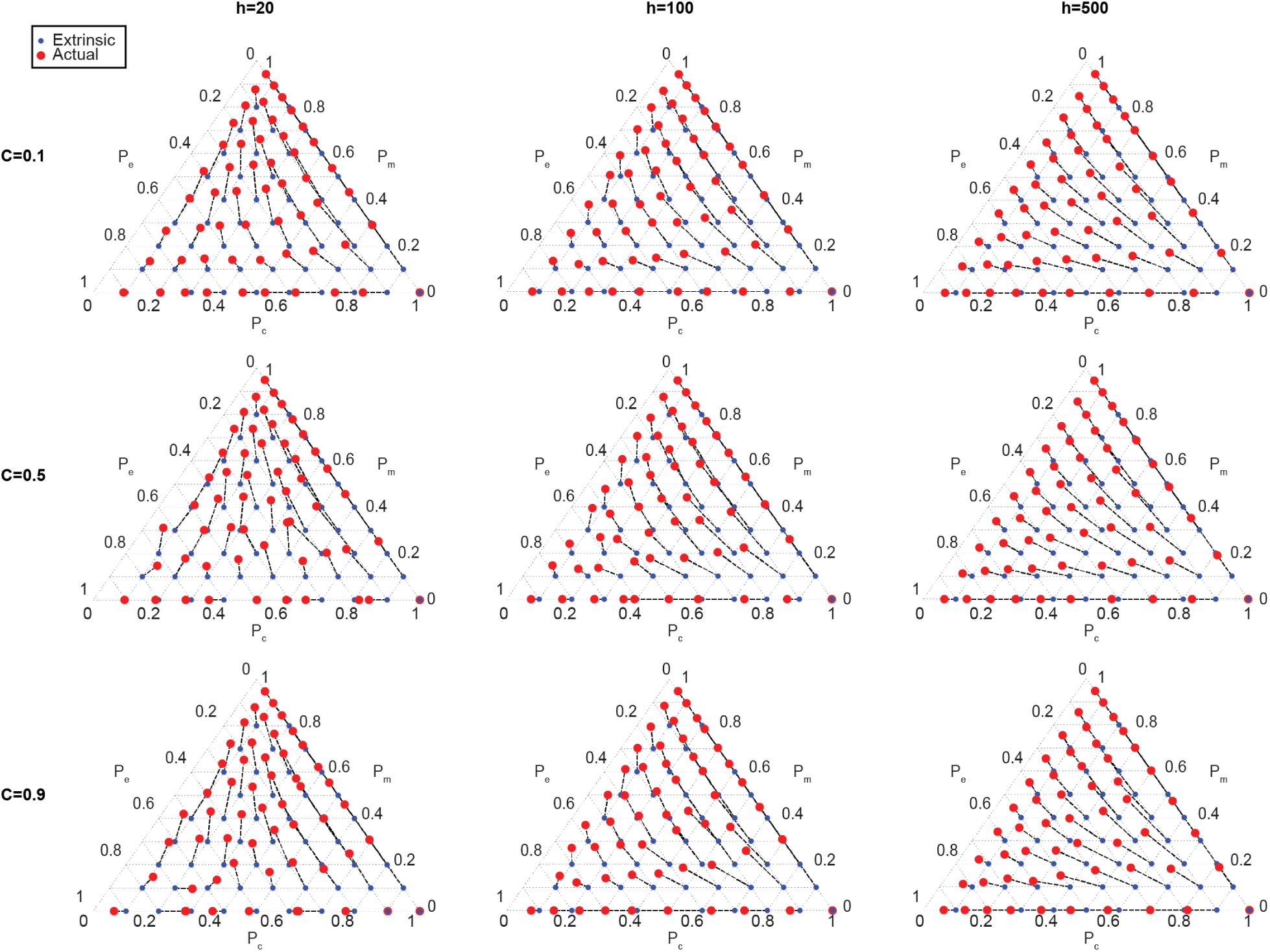
The extrinsic (blue points) and the actual (red points) interaction proportions of species that go extinct, for the UIM for varying *C* and *h*. Dotted lines connect points from the same community.

**Supplementary Figure 18:**
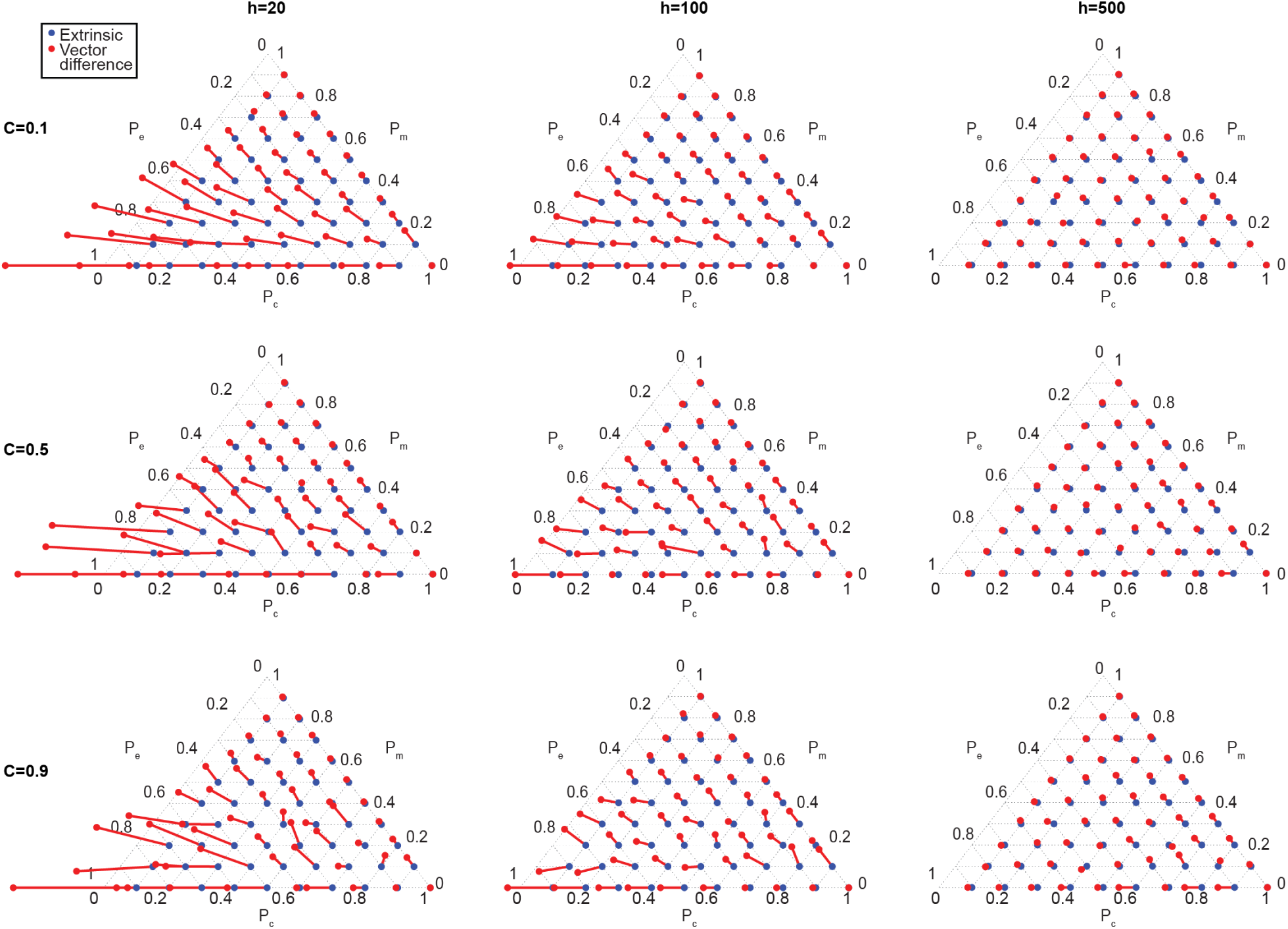
For the UIM, the vector difference between species that go extinct and species that invade the community are plotted as red lines with red points at the end. The vectors start from the extrinsic proportions (blue points) and are defined as the interaction types of species that go extinct minus the interaction types of invaders.

**Supplementary Figure 19:**
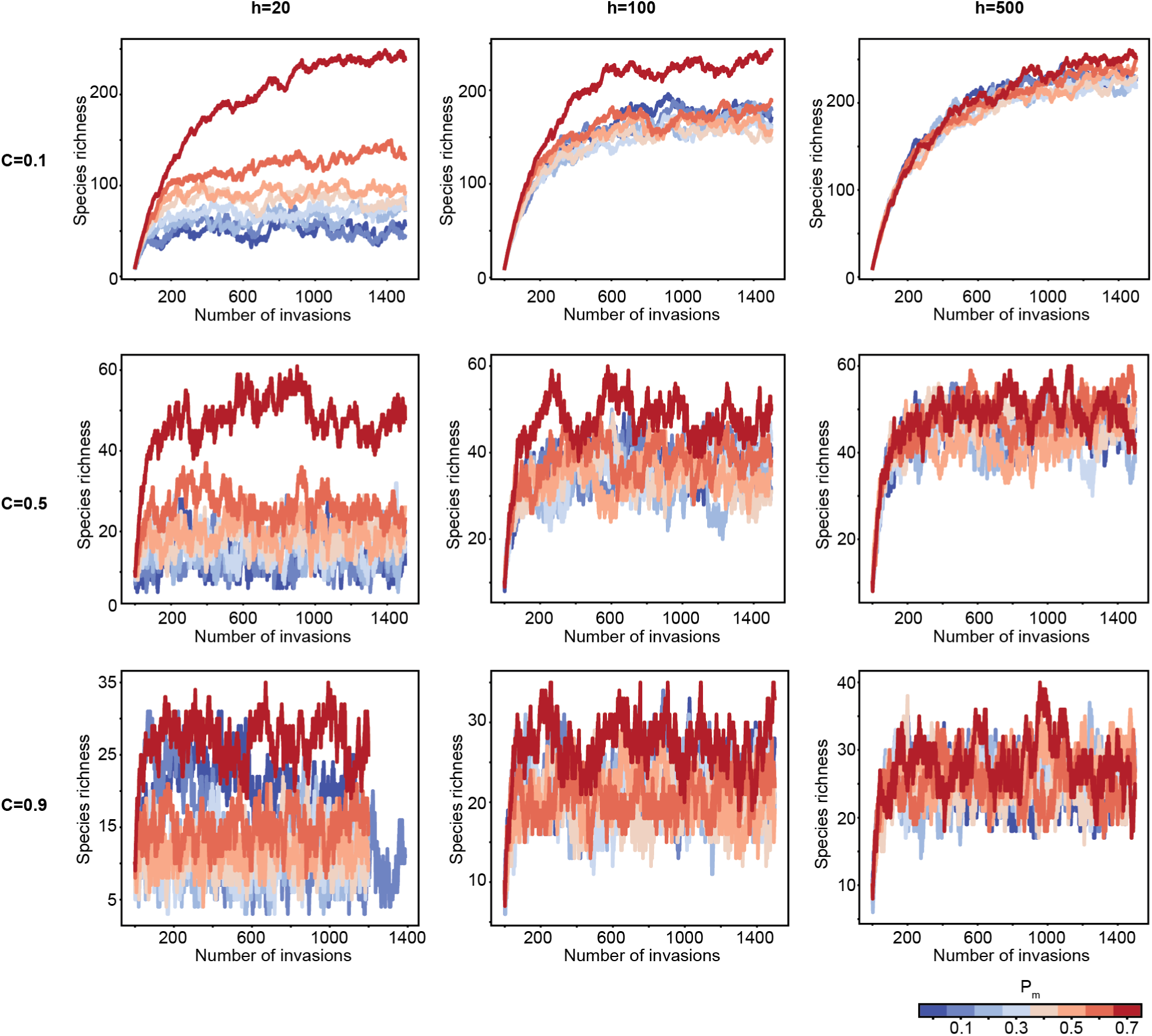
For the IIM, the mixture of interaction types determines the species richness at which communities saturate, for various *C* and *h*. Species richness in representative communities with *P_c_* = 0.3 and varying levels of *P_m_*, represented by the color. The colors in all plots correspond to the color bar shown. Each line corresponds to a community over 1500 species invasions that occur at equilibria. The same trend is observed for all *P_c_ >* 0.

**Supplementary Figure 20:**
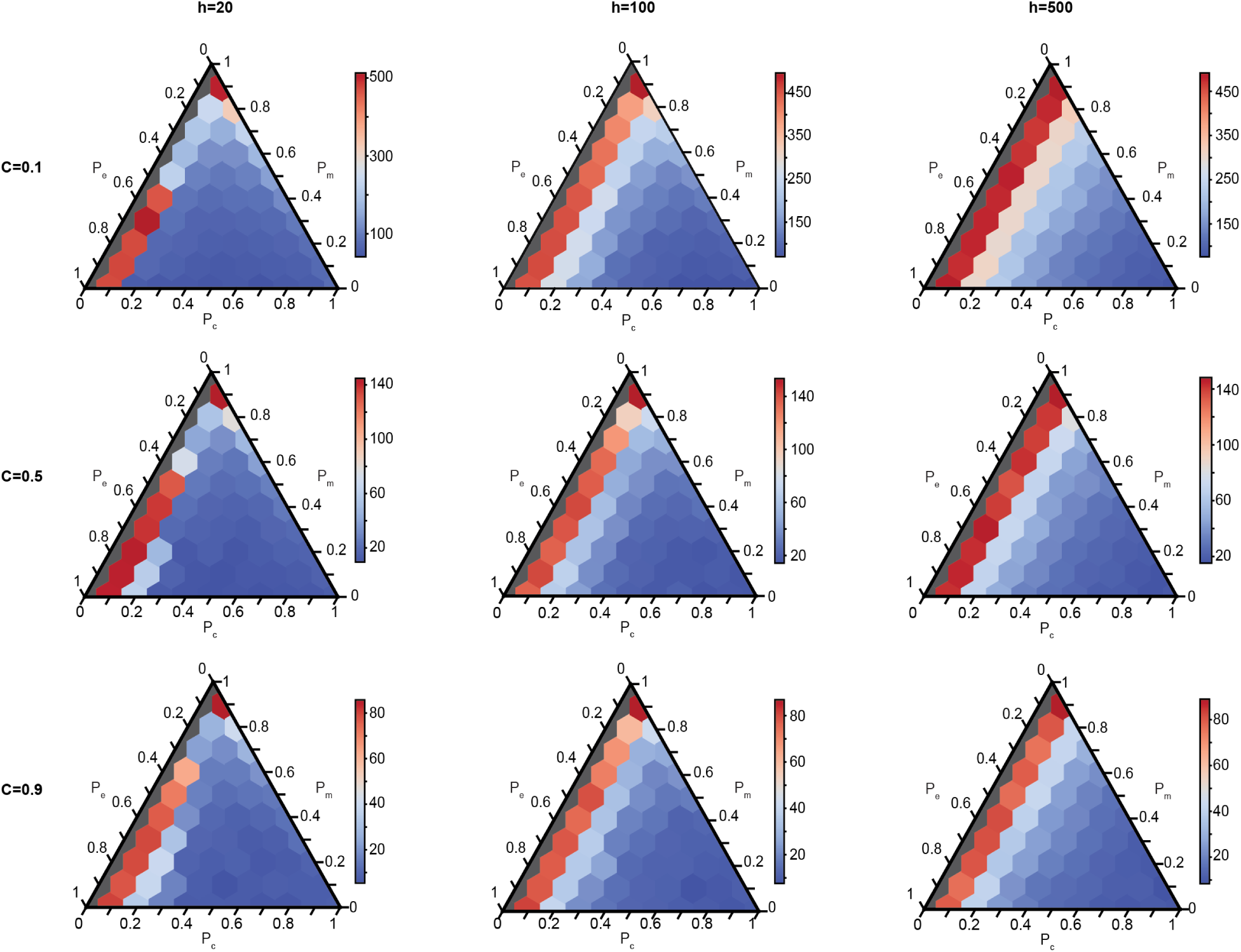
For the IIM, the steady state species richness of different communities with varying proportions of competition, mutualism, and exploitation, and varying *C* and *h*. The color represents the species richness, and each plot has its own color scale. Gray communities have no competition and are not considered since they are biologically unrealistic.

**Supplementary Figure 21:**
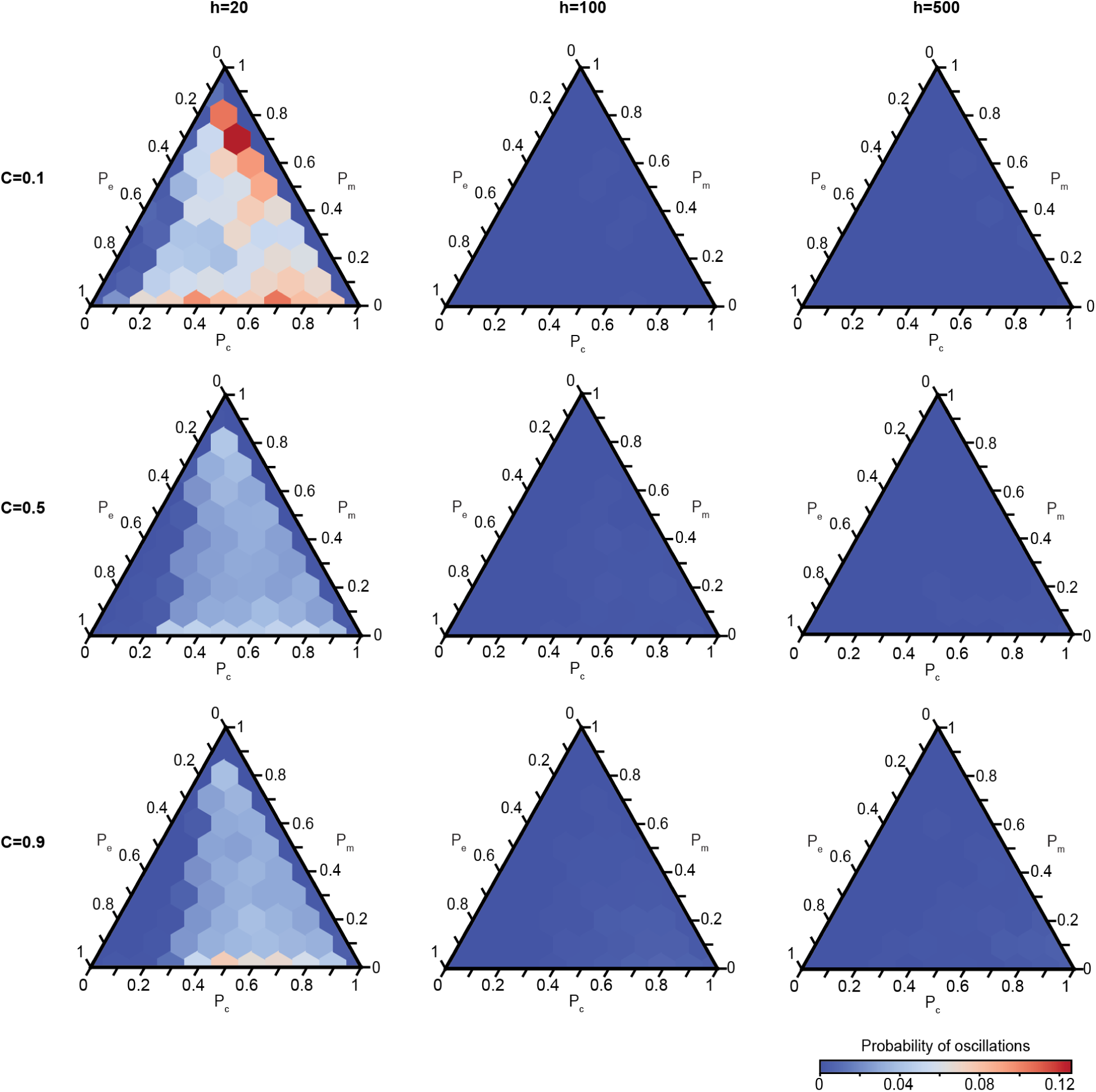
For the IIM, the probability of oscillations throughout the entire community history, for all combinations of interaction types, *C*, and *h*. The color represents the probability and all plots use the same scale, following the color bar shown.

**Supplementary Figure 22:**
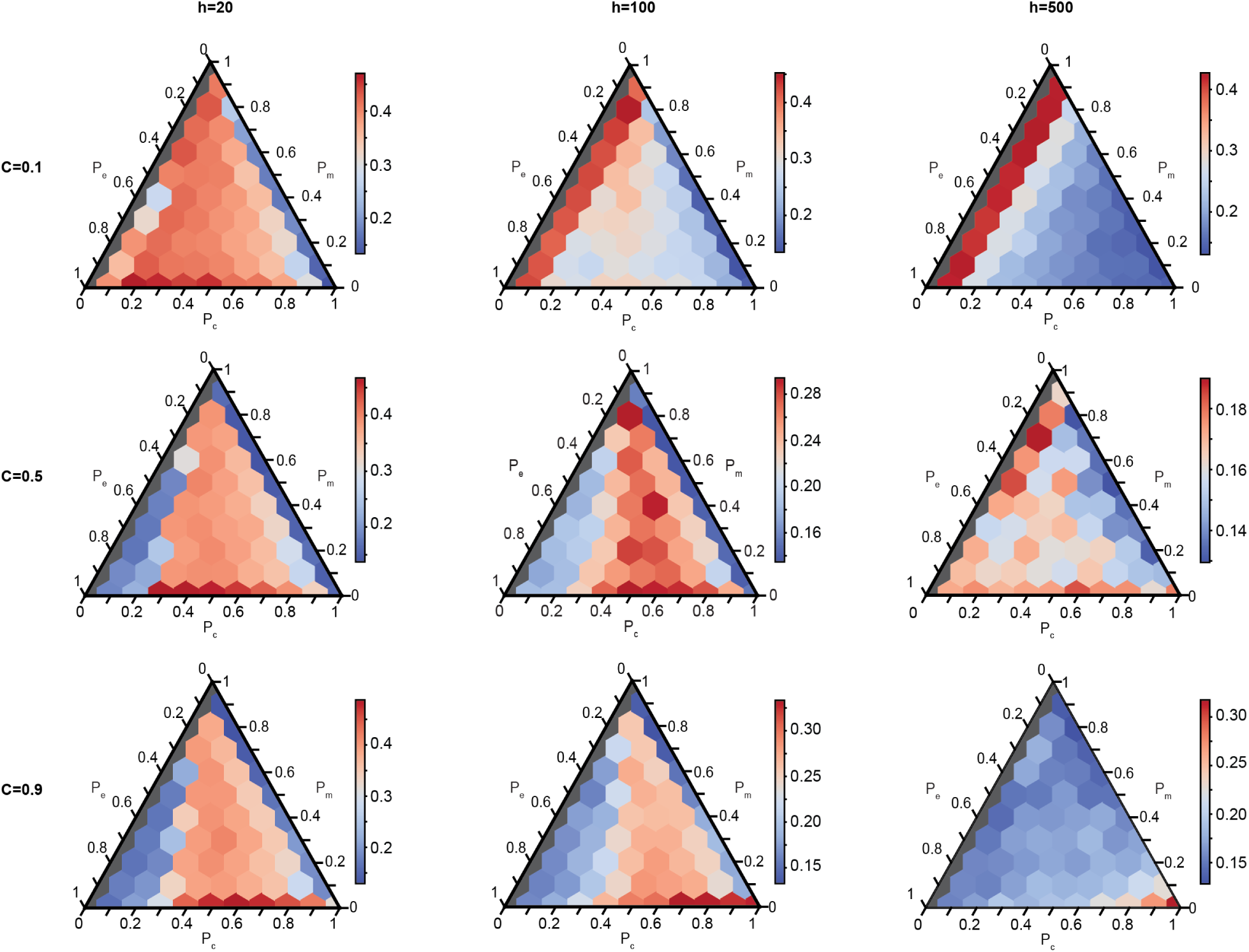
External stability throughout the entire community history for the IIM for varying *C*, *h*, and proportion of interaction types. Colors indicate the probability of invasion and each plot has its own color scale. Gray communities have no competition and are not considered.

**Supplementary Figure 23:**
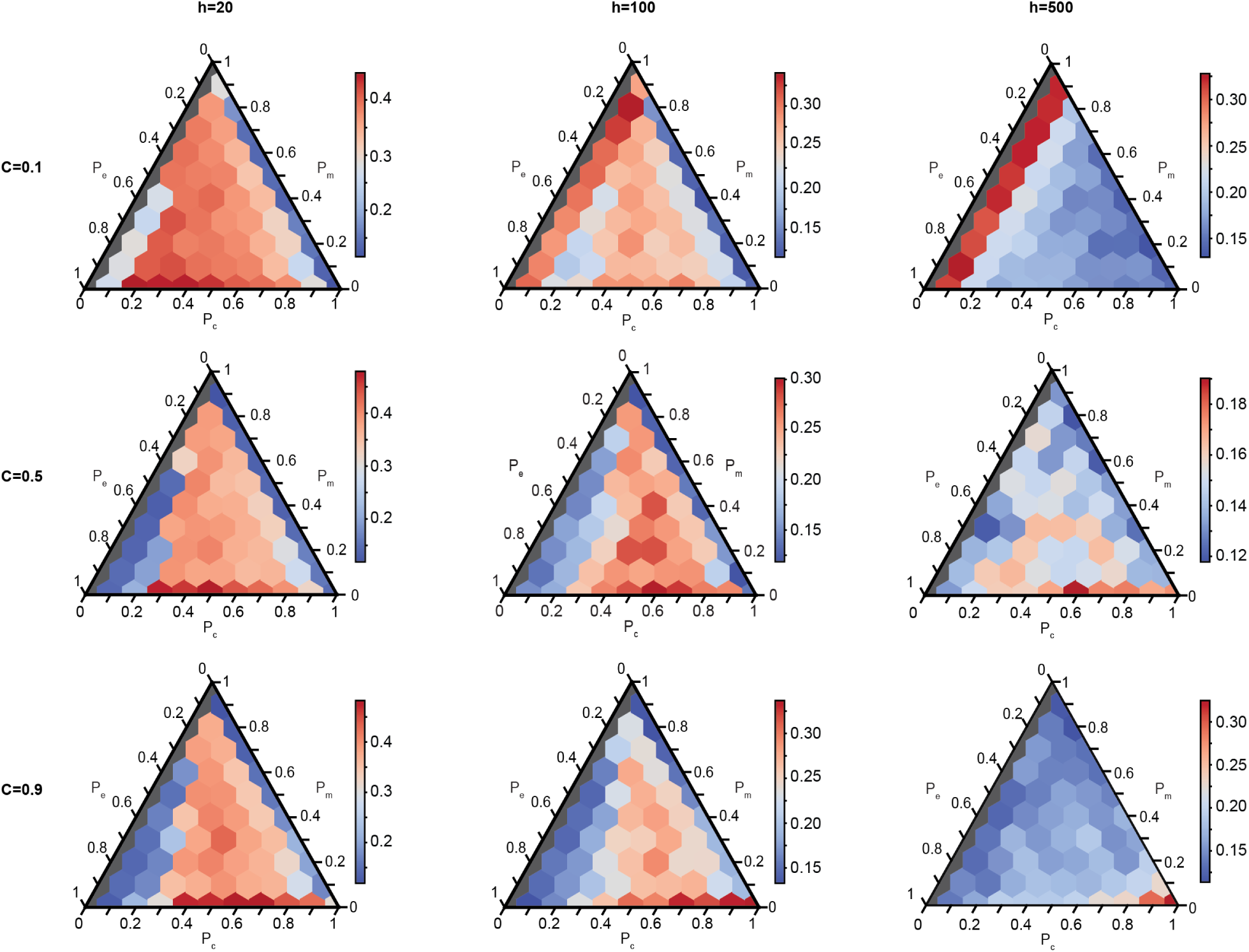
External stability after species richness has converged to the steady state for the IIM with varying *C*, *h*, and proportion of interaction types. Colors indicate the probability of invasion and each plot has its own color scale. Gray communities have no competition and are not considered.

**Supplementary Figure 24:**
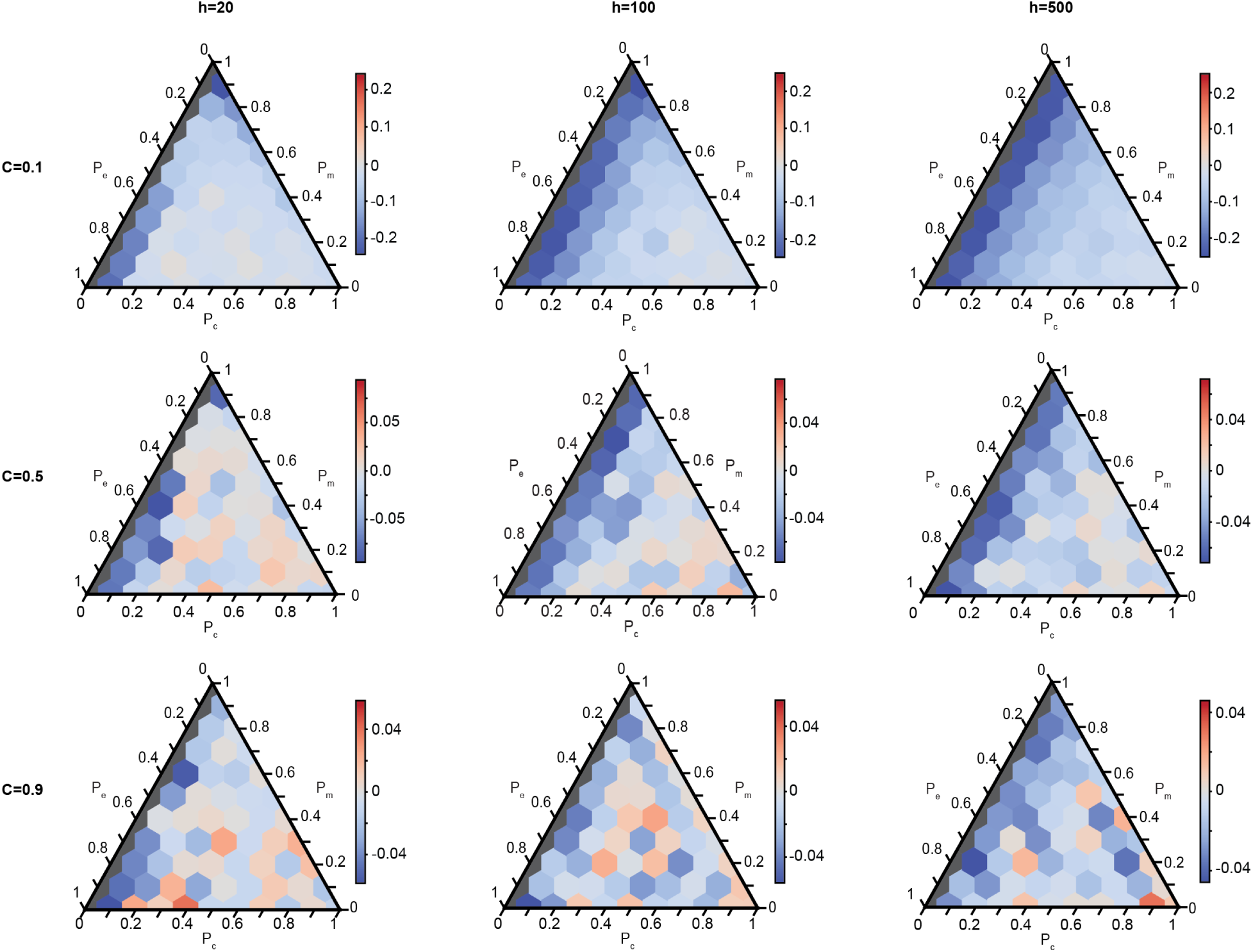
The difference in probability of invasion between the transient state and the steady state for the IIM for varying *C*, *h*, and proportion of interaction types. Colors indicate the change in the probability of invasion and each plot has its own color scale. Gray communities have no competition and are not considered.

**Supplementary Figure 25:**
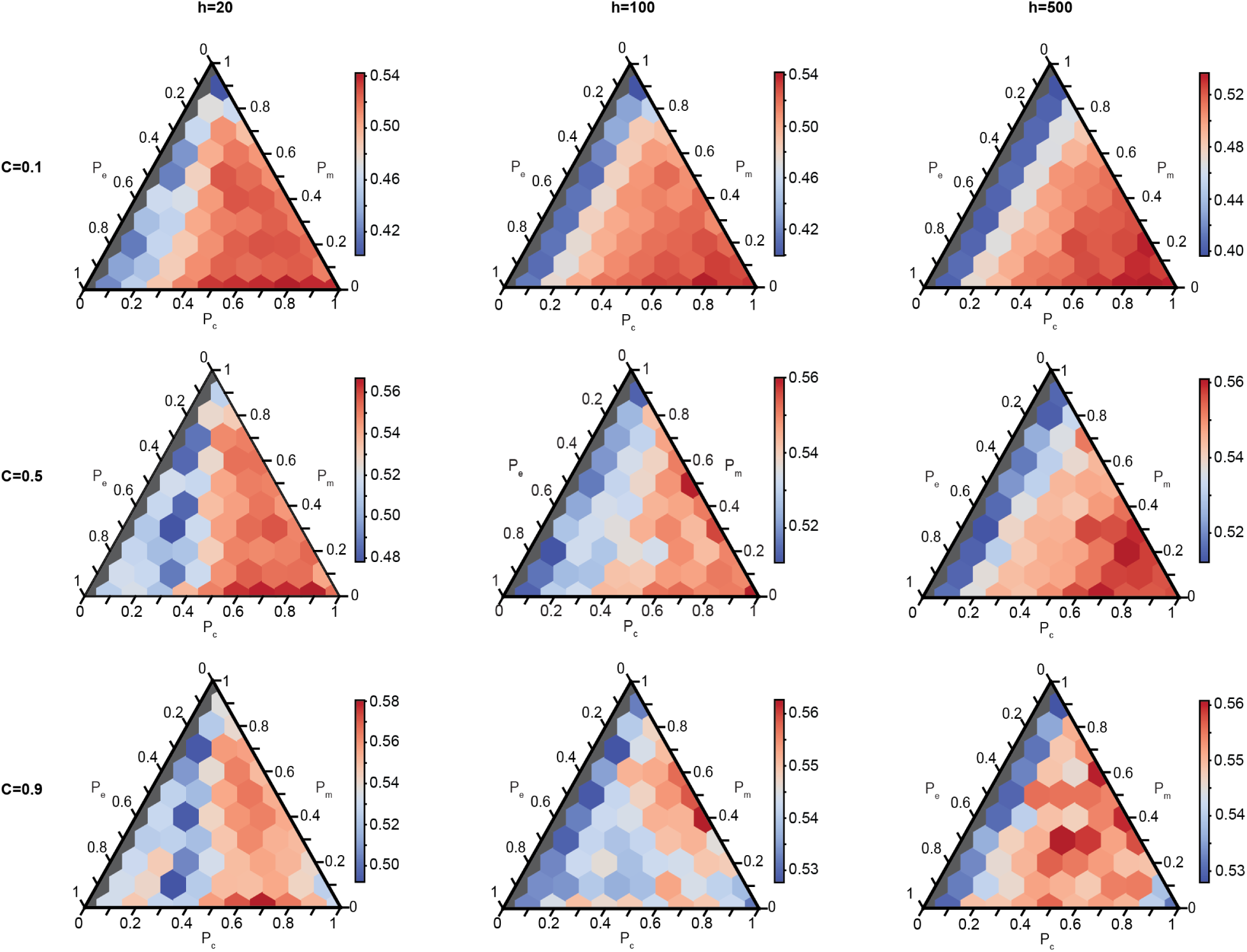
Probability of extinction throughout the entire community history for the IIM for varying *C*, *h*, and proportion of interaction types. Colors indicate the probability of extinction and each plot has its own color scale. Gray communities have no competition and are not considered.

**Supplementary Figure 26:**
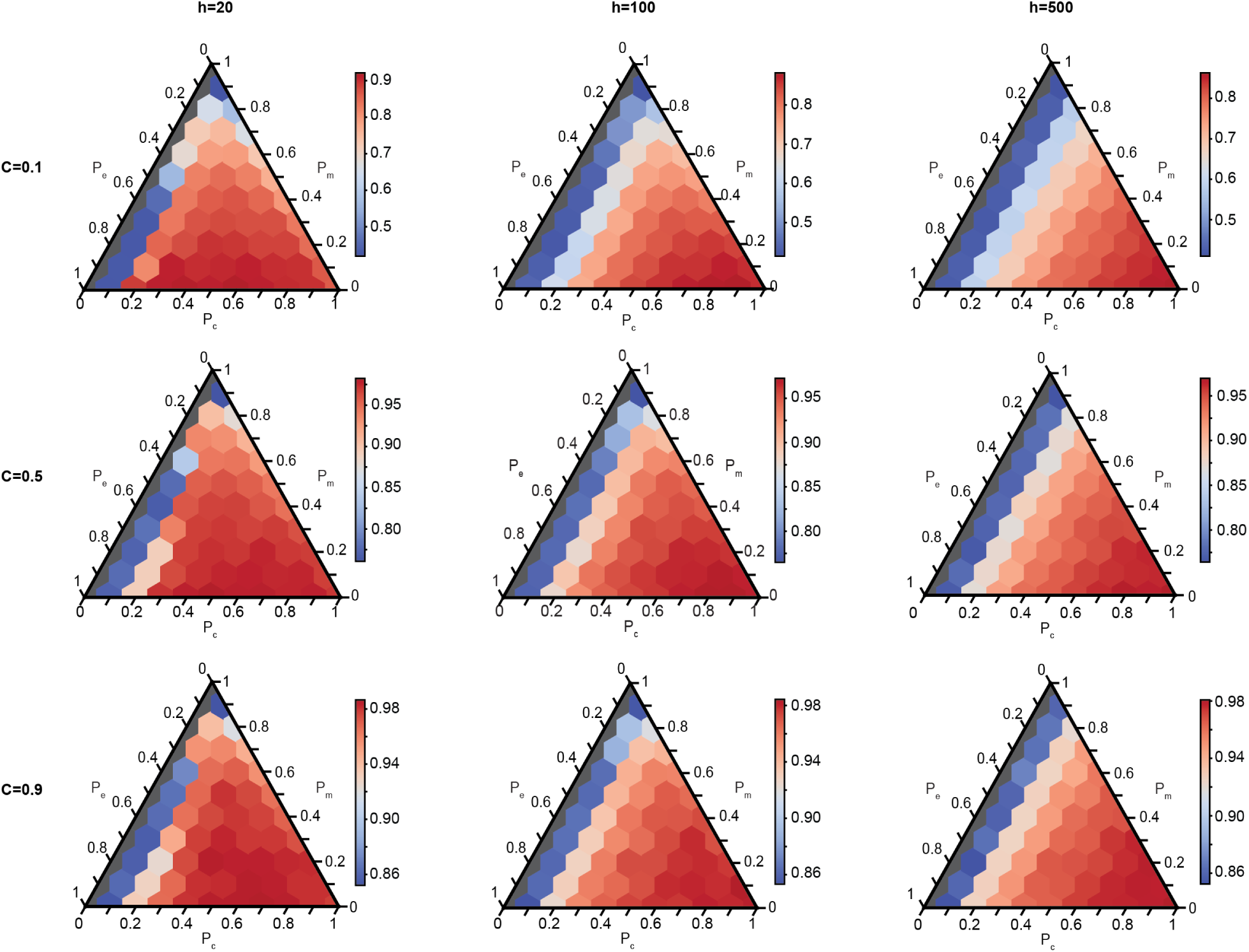
Probability of extinction after species richness has converged to the steady state for the IIM for varying *C*, *h*, and proportion of interaction types. Colors indicate the probability of extinction and each plot has its own color scale. Gray communities have no competition and are not considered.

**Supplementary Figure 27:**
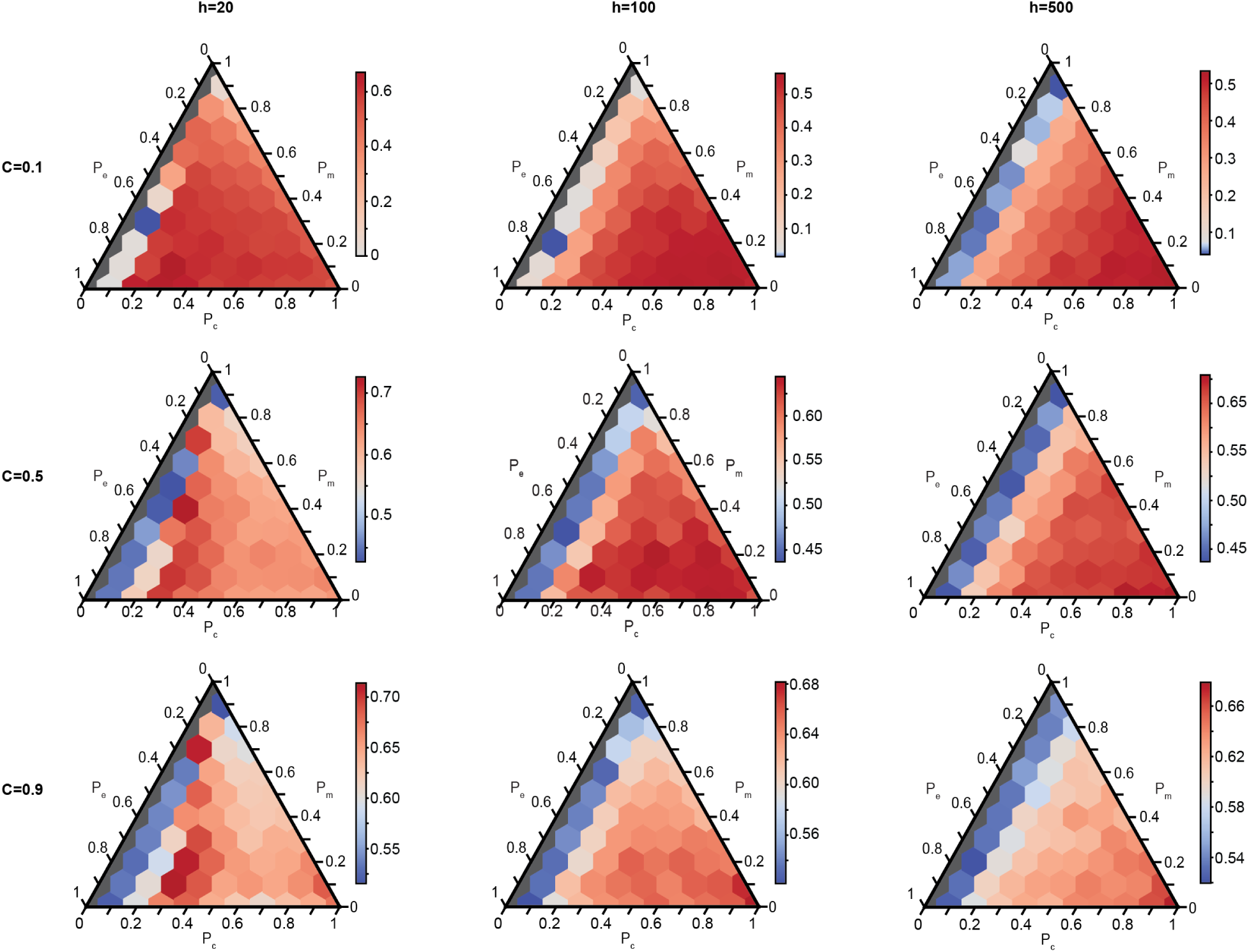
The difference in probability of extinction between the transient state and the steady state for the IIM for varying *C*, *h*, and proportion of interaction types. Colors indicate the change in the probability of extinction and each plot has its own color scale. Gray communities have no competition and are not considered.

**Supplementary Figure 28:**
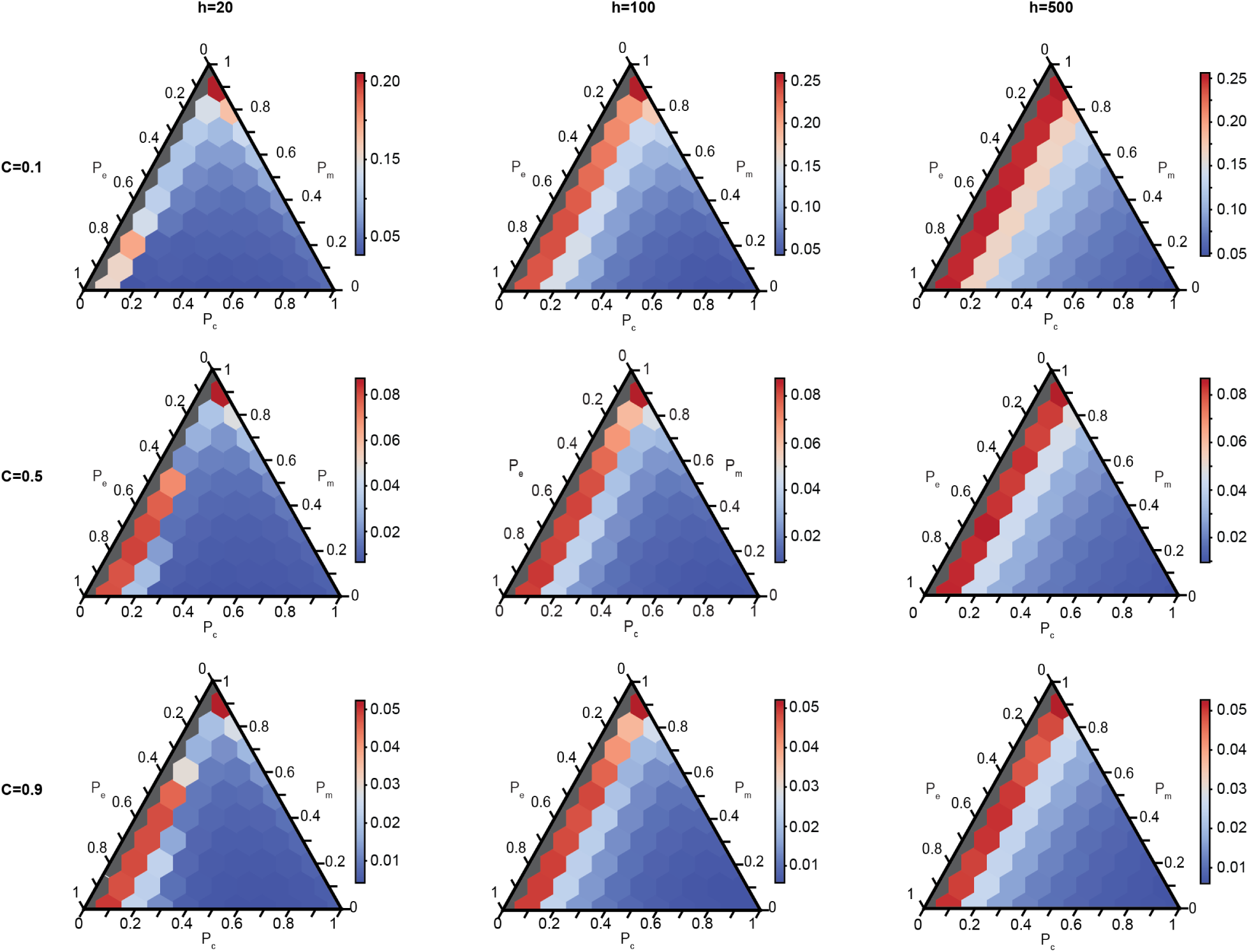
The mean relative persistence of all species that existed within a community for the IIM for varying *C*, *h*, and proportion of interaction types. The color represents the mean relative persistence, and each plot has its own color scale. Gray communities have no competition and are not considered.

**Supplementary Figure 29:**
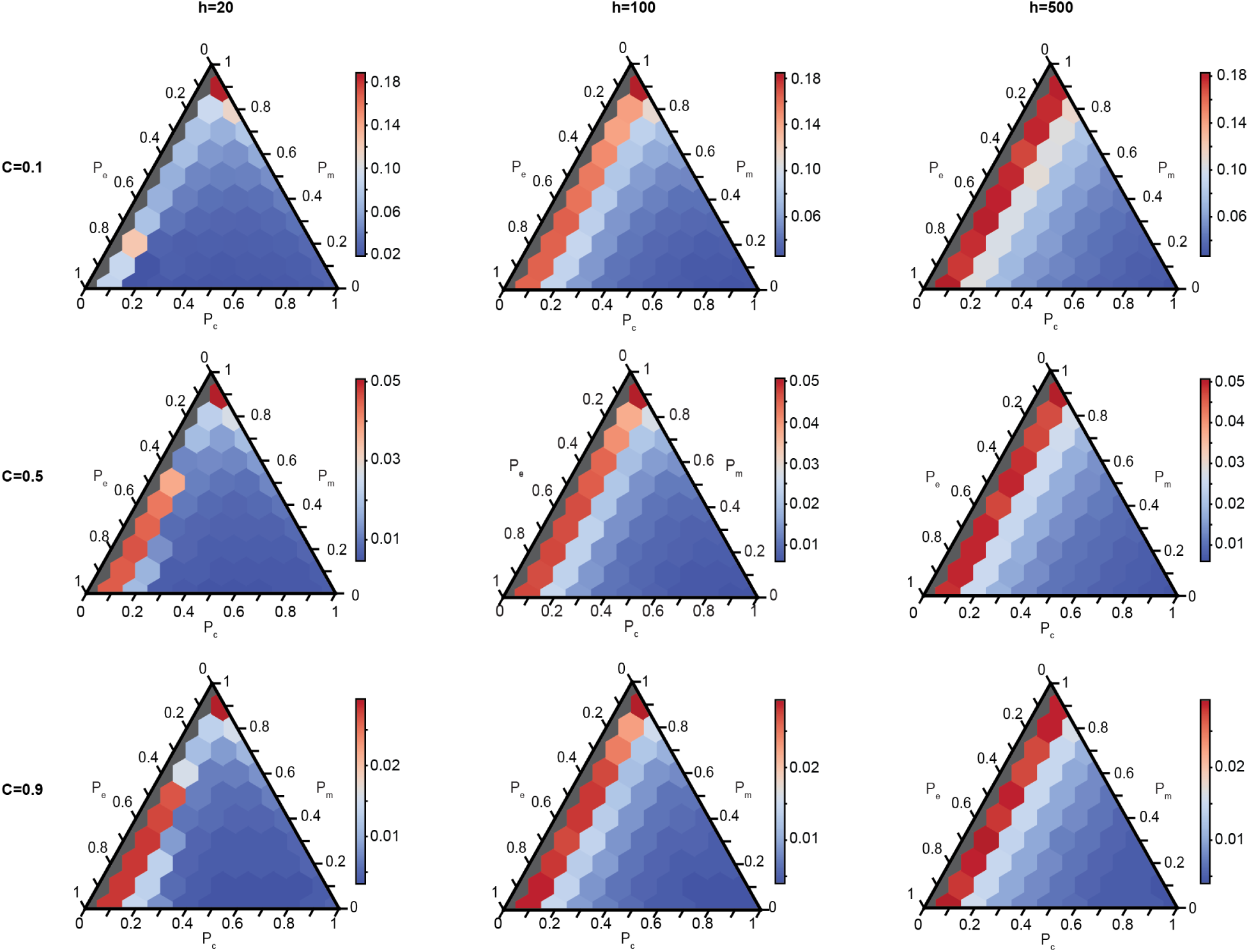
The median relative persistence of all species that existed within a community for the IIM for varying *C*, *h*, and proportion of interaction types. The color represents the median relative persistence, and each plot has its own color scale. Gray communities have no competition and are not considered.

**Supplementary Figure 30:**
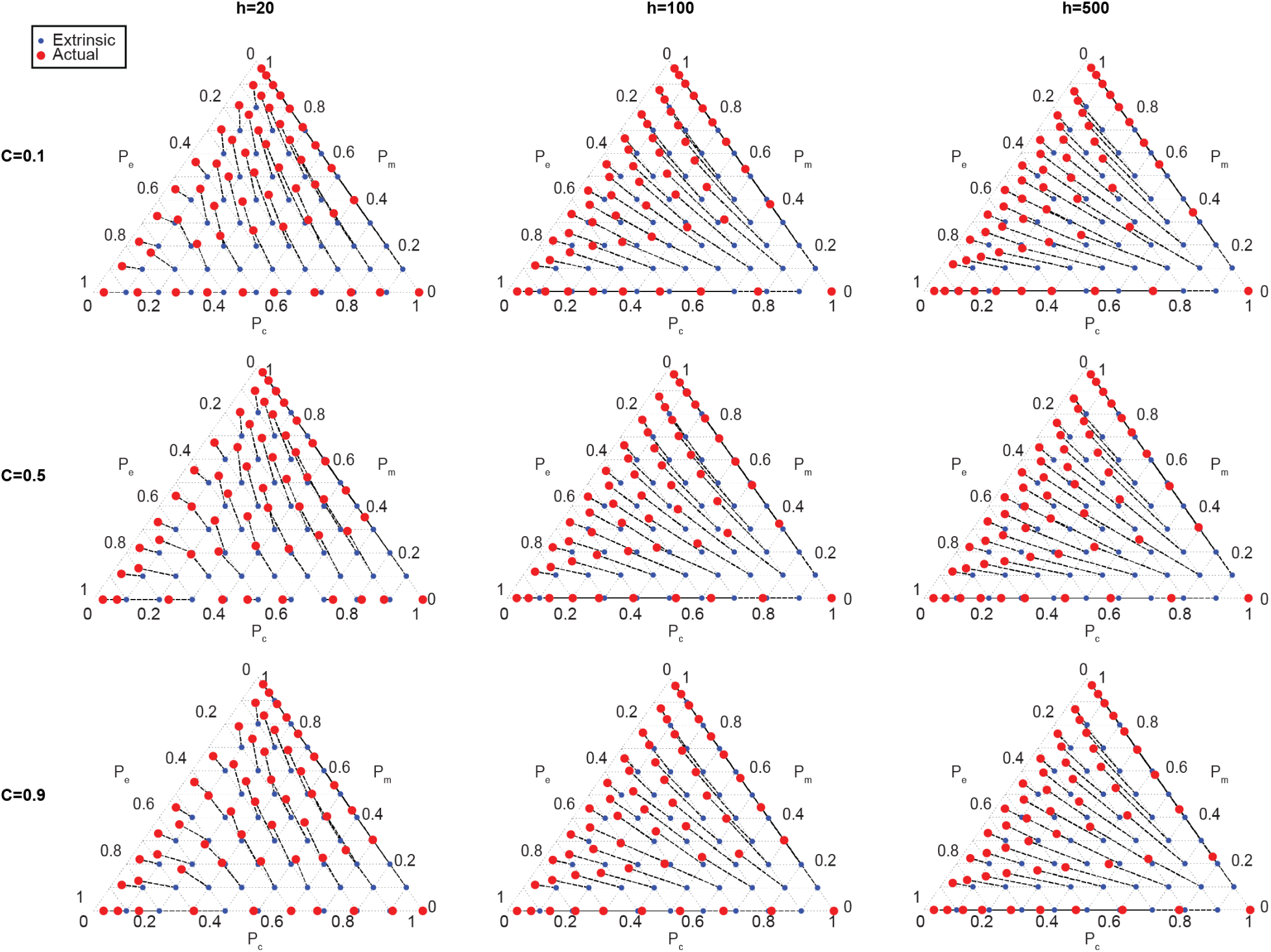
The extrinsic (blue points) and the actual (red circles) interaction proportions of all species within each community for the IIM for varying *C* and *h*. Dotted lines connect points from the same community, and can be interpreted as the direction of selection.

**Supplementary Figure 31:**
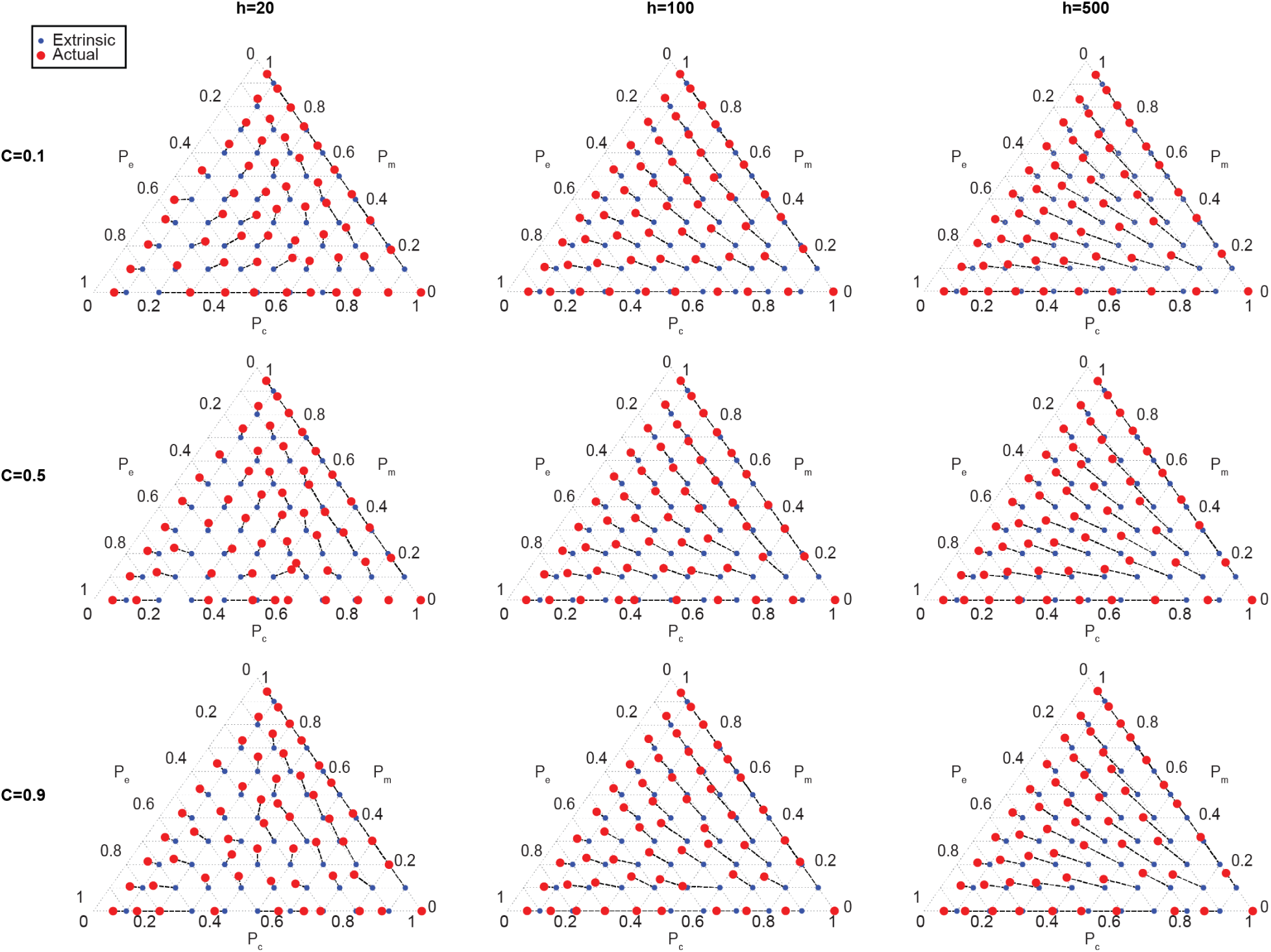
The extrinsic (blue points) and the actual (red circles) interaction proportions of species that successfully invade each community for the IIM for varying *C* and *h*. Dotted lines connect points from the same community, and can be interpreted as the direction of selection.

**Supplementary Figure 32:**
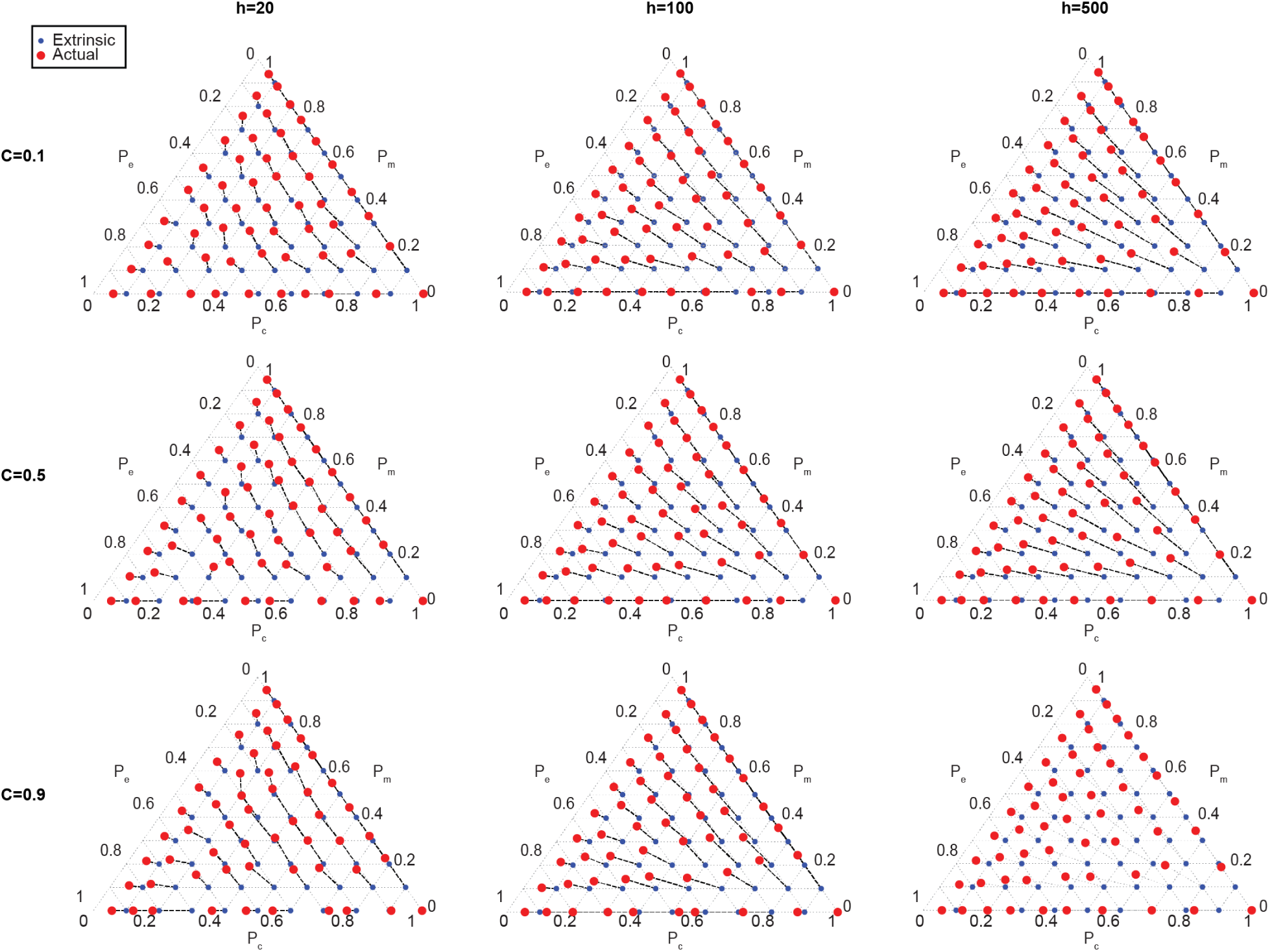
The extrinsic (blue points) and the actual (red points) interaction proportions of species that go extinct, for the IIM for varying *C* and *h*. Dotted lines connect points from the same community.

**Supplementary Figure 33:**
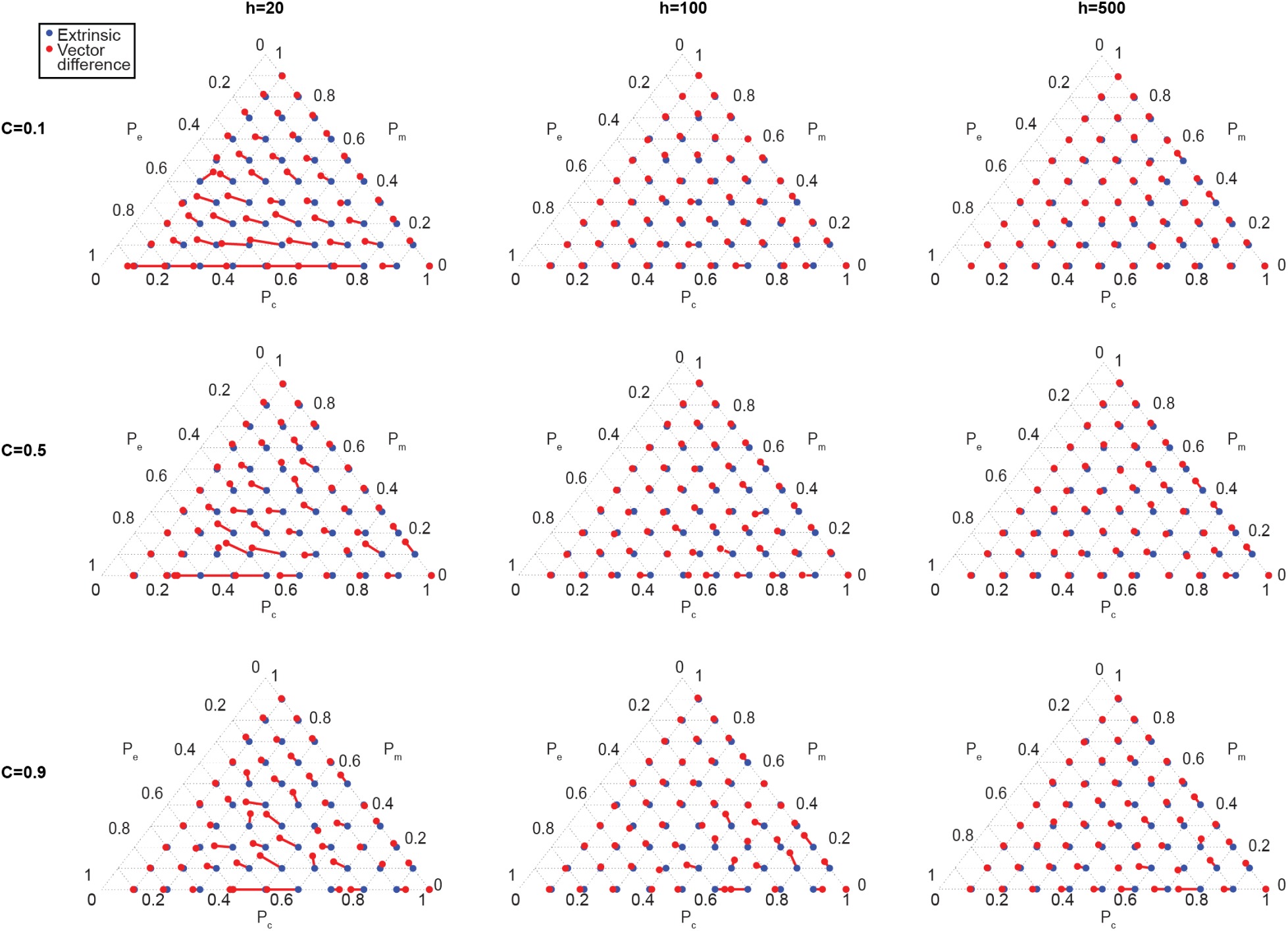
For the IIM, the vector difference between species that go extinct and species that invade the community are plotted as red lines with red points at the end. The vectors start from the extrinsic proportions (blue points) and are defined as the interaction types of species that go extinct minus the interaction types of invaders.

